# Imp and Syp *in vivo* temporal RNA interactomes uncover networks of temporal regulators of *Drosophila* brain development

**DOI:** 10.1101/2024.06.30.600407

**Authors:** Jeffrey Y Lee, Niles Huang, Tamsin J Samuels, Ilan Davis

## Abstract

Temporal patterning of neural progenitors is an evolutionarily conserved mechanism generating neural diversity. In *Drosophila*, post-embryonic neurogenesis requires the RNA-binding proteins (RBPs) Imp/IGF2BP and Syp/SYNCRIP. However, how they co-achieve their function is not well understood. Here, we elucidate the *in vivo* temporal RNA interactome landscapes of Imp and Syp during larval brain development. Imp and Syp bind a highly overlapping set of conserved mRNAs encoding proteins involved in neurodevelopment. We identify transcripts differentially occupied by Imp/Syp over time, featuring a network of known and novel candidate temporal regulators that are post-transcriptionally regulated by Imp/Syp. Furthermore, the physical and co-evolutionary relationships between Imp and Syp binding sites reveal a combinatorial, rather than competitive, mode of molecular interplay. Our study establishes a new *in vivo* framework for dissecting the temporal co-regulation of RBP networks as well as providing a resource for understanding neural fate specification.

## Introduction

Understanding how a small population of neural stem cells (NSCs) generate the complex brain is a key question in developmental biology with significant biomedical implications, as many neurological disorders stem from aberrant neurodevelopment (Pilaz and Silver, 2015; Yano et al., 2016). *Drosophila melanogaster* serves as an excellent model for brain development studies, as it shares many key features with mammalian neurogenesis, including NSC delamination (Skeath and Thor, 2003), the balance between NSC self-renewal and differentiation (Doe, 2008; Homem and Knoblich, 2012), and tumourigenesis upon uncontrolled proliferation of progenitors (Caussinus and Hirth, 2007; Homem et al., 2015; Maurange and Gould, 2005). Importantly, the principle of increasing neural diversity through temporally expressed factors is also conserved in *Drosophila*, making it an ideal model to study this mechanism (Doe, 2017, 2006; Isshiki et al., 2001; Konstantinides et al., 2022; Maurange et al., 2008; Syed et al., 2017). In this process, molecular signatures of upstream NSCs underpin the morphological diversity of immature neurons (Kepecs and Fishell, 2014). Therefore, elucidating the complex landscapes of gene expression regulation is a critical step in understanding specification of neuronal identity.

Temporal patterning of neuroblasts (NBs, *Drosophila* NSCs) has been most extensively studied in the embryonic ventral nerve cord (VNC) where cascades of temporal transcription factors (TFs) generate birth-order-dependent neural diversity (Isshiki et al., 2001). This involves cross-regulatory control between temporal TFs that drives fate transitions (Doe, 2017). Recently, post-transcriptional regulation via RNA stability, translation and localisation has emerged as another layer of diversity generation. RNA-binding proteins (RBPs) are the key mediators of post-transcriptional regulation, adding complexity and precision to gene expression output (Franks et al., 2017; Pilaz and Silver, 2015). Temporal patterning during larval and pupal stages features opposing gradients of conserved RBPs, IGF2 mRNA-binding protein (Imp/IGF2BP) and Syncrip (Syp/SYNCRIP), which negatively regulate each other (Liu et al., 2015; Ren et al., 2017). Unlike short-range transitions driven by embryonic temporal TFs, the gradually changing levels of Imp and Syp in post-embryonic NB lineages produce a greater number and diversity of cell types (Islam and Erclik, 2022). Thus, the reciprocity between DNA and RNA-level regulations is likely to be critical to the fate specification programme, accommodating the demands of complex adult functions.

Imp and Syp levels are known to influence specification of the mushroom body (Liu et al., 2020, 2015), motor neurons (Guan et al., 2022), the central complex (Hamid et al., 2023), and the visual system (Arain et al., 2022). Beyond fate patterning, they also regulate NB quiescence exit (Munroe et al., 2022), NB growth and self-renewal (Hailstone et al., 2020; Landskron et al., 2018; Samuels et al., 2020b), NB decommissioning (Pahl et al., 2019; C. P. Yang et al., 2017), synaptic transmission (Boylan et al., 2008; Halstead et al., 2014; Hansen et al., 2015; Medioni et al., 2014; Titlow et al., 2020), and fly behaviour (Hamid et al., 2023; Samuels et al., 2020a). Furthermore, the protracted nature of Imp and Syp gradients allows integration of extrinsic signals into the intrinsic patterning programme (e.g. steroid hormone, Activin and Notch signalling) (Branham et al., 2024; Rossi and Desplan, 2020; Syed et al., 2017). Although some individual downstream targets of Imp and Syp have been characterised in the studies cited above, the complete RNA interactomes and their temporal variations remain unexplored. Recent single-cell RNA-seq (scRNA-seq) studies have revealed RNAs displaying cell-type and development-specific expressions in the larval brain (Corrales et al., 2022; Dillon et al., 2022; Janssens et al., 2022; Ravenscroft et al., 2020). Therefore, it is pertinent to identify which of these transcripts are targeted by Imp and Syp, as they may represent potential downstream effectors of fate patterning.

Here, we identify the RNA interactomes of Imp and Syp across early and late stages of the developing larval brain. By adapting iCLIP (individual nucleotide-resolution UV crosslinking immunoprecipitation) in larval tissues, we uncover *in vivo* binding sites of Imp and Syp at an unprecedented resolution. Our dataset reveals highly overlapping Imp and Syp targets enriched for gene expression regulators with diverse neurodevelopmental roles. Leveraging the temporal aspect of our dataset, we identify transcripts that are differentially occupied by Imp and Syp across development, uncovering a complex network of temporal regulators or genes with as-yet-unknown fate patterning functions acting downstream of Imp and Syp. Furthermore, we show key examples of Imp/Syp-mediated post-transcriptional regulation of downstream targets that influence early or late stage neuronal identity. Finally, we profile physical and evolutionary relationships between Imp and Syp binding sites to provide insights into the molecular interplay underpinning their regulatory cassette. Our comprehensive analysis of the downstream landscapes of Imp and Syp can now serve the community in finely dissecting regulatory mechanisms of temporal cell fate specification, as well as providing a resource for studying regulatory interplay between RBPs.

## Results

### Identification of Imp and Syp targets across larval brain development

To identify *in vivo* RNA interactomes of Imp and Syp, we adapted iCLIP2 (Buchbender et al., 2019) for *Drosophila* larval tissues and mapped their transcriptome-wide binding sites. We chose three post-embryonic developmental time points: L1, L2 and L3 (24h, 48h and 96h after larval hatching (ALH)), representing different stages of graded Imp and Syp protein expression levels (Liu et al., 2015; Syed et al., 2017) (Figure 1A and Figure S1A). For the L1 stage, Imp is highly enriched in the L1 CNS (Figure 1A), therefore we immunoprecipitated endogenously tagged Imp::FLAG (*Imp::GFSTF*) from frozen whole-larval powder after UV-crosslinking. At the L2 stage, both Imp and Syp proteins are highly enriched in the larval CNS, however we used severed larval heads to immunoprecipitate both Imp::GFSTF and Syp to avoid contamination from testes-expressed Syp RNP (McDermott et al., 2012). For the L3 stage, we individually dissected larval brains and used high-affinity GFP-trap antibody against endogenously tagged Syp::eGFP (Titlow et al., 2020) to maximise RNP recovery. Additionally, we prepared corresponding size-matched input libraries (SMInput), which represents the control background (Figure S1B) (Van Nostrand et al., 2016). Principal component analysis (PCA) of the sequenced libraries demonstrated clear separation between iCLIP and SMInput libraries and close clustering of biological replicates (Figure S1C-D), confirming the reproducibility of our experiments. Furthermore, iCLIP reads of Imp and Syp predominantly mapped to 3’UTR or non-coding RNA (ncRNA) features, while SMInput libraries were over-represented in CDS, intronic, and intergenic elements, confirming the specificity of the immunoprecipitated RNA fragments (Figure S1E).

**Figure 1.**
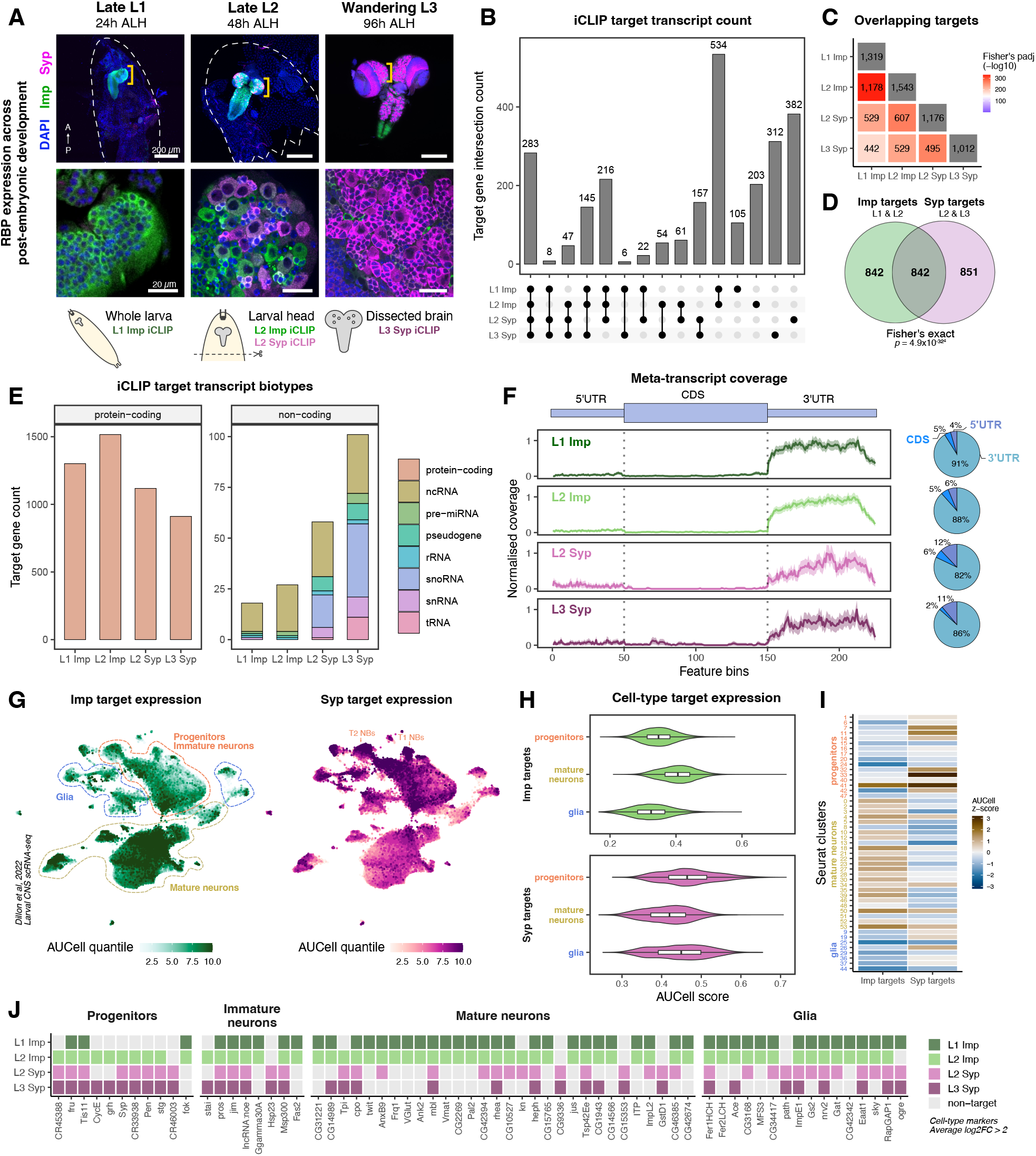
Identification of *in vivo* RNA targets of Imp and Syp in larval brains. **(A)** Temporal expression pattern of Imp and Syp protein in *Drosophila* larval nervous system. Dotted regions indicate biosamples collected for individual nucleotide-resolution UV crosslinking and immunoprecipitation (iCLIP). **(B)** Intersections between Imp and Syp iCLIP target transcripts. **(C)** The statistical significance of target transcript overlap between each iCLIP library, assessed using the hypergeometric test. **(D)** The statistical significance of target transcript overlap between Imp targets (L1 and L2 combined) and Syp targets (L2 and L3 combined). **(E)**Relative proportions of Imp and Syp iCLIP targets by their transcript biotypes. **(F)** Meta-transcript coverage of Imp and Syp binding sites on protein-coding genes. Each transcript was represented by its longest coding sequence (CDS) and untranslated region (UTR) isoforms for the coverage calculation. Pie charts represent distribution of Imp and Syp binding sites in CDS, 5’UTR and 3’UTR regions. **(G)** Area Under Curve Cell (AUCell) scores of Imp and Syp targets across the larval brain single-cell RNA-seq atlas (Dillon et al., 2022), which represents relative RBP target expression levels (see Methods). Progenitors/immature neurons, mature neurons and glial cell types identified from the original publication are indicated. **(H)** Comparison of AUCell scores of Imp and Syp targets between progenitors, mature neurons or glia cell-types. **(I)** Heatmap of average AUCell scores of Imp and Syp targets in different cell types. The Seurat clusters and their broad cell types (progenitors, immature neurons, mature neurons, glia) are indicated. **(J)** Cell type-specific transcripts that are Imp or Syp targets. Only the transcripts that were identified from at least one iCLIP library are shown.

We determined nucleotide-resolution binding sites of Imp and Syp by clustering significant crosslinks (FDR < 0.01) and filtering for enrichment over SMInput samples (Lee and Ule, 2018; Sahadevan et al., 2022). This analysis identified 1,000 - 1,500 transcriptome-wide targets of Imp and Syp, harbouring significant RBP binding sites (Figure 1B and Supplementary Table 1). Of note, the lack of correlation between iCLIP crosslink reads and transcript expression levels suggests that Imp and Syp bind to their targets with specificity, rather than passively associating with abundant transcripts (Figure S1F). Interestingly, we found a significant overlap between Imp and Syp targets (Figure 1C-D), indicating a large cohort of transcripts regulated either competitively or cooperatively. Both RBPs primarily targeted protein coding transcripts (Figure 1E) with a strong preference to 3’UTRs (Figure 1F), though an increasing proportion of crosslinks to ncRNAs was noted in L2 and L3 libraries (Figure 1E). Since Imp and Syp are highly conserved RBPs across the animal kingdom, we asked whether their RNA targets are also conserved. To this end, we compared our dataset with the RNA targets of human IMP1-3 in pluripotent stem cells (Conway et al., 2016) and murine Syncrip/hnRNPR in neural cells (Briese et al., 2018; Khudayberdiev et al., 2021). Within the genes that had high-confidence orthologues (DIOPT score > 8), we found a significant degree of conservation of RBP targets between fly and mammals: 76% (871/1144) for Imp and 55% (623/1134) for Syp (Supplementary Table 1). This result suggests evolutionarily conserved RNA-binding specificities of Imp and Syp.

Next, we examined the general expression patterns of Imp and Syp targets using recently published larval brain single-cell RNA-seq (scRNA-seq) atlases (Aibar et al., 2017; Corrales et al., 2022; Dillon et al., 2022; Ravenscroft et al., 2020). Grouping cell-type clusters into progenitors, immature neurons and mature neurons, we found a general trend where Imp targets were more highly expressed in mature neurons compared to progenitor cells, while Syp targets were more highly expressed in progenitor cells (Figure 1G-I and Figure S1G-H). Immature neurons showed intermediate levels of Imp and Syp target expressions. This observation correlates well with spatial RBP expression patterns, as Imp is expressed in early-born functional neurons and Syp in progenitors later during development (Allen et al., 2020; Syed et al., 2017). Nevertheless, we found Imp and Syp bind a wide range of cell-type marker genes across the neuronal differentiation trajectory as well as glia, indicating their tissue-wide activity (Figure 1J). Significant binding sites of Imp and Syp were also found in transcripts specifically expressed in rare NB cell types (e.g. *CycE*, *lncRNA::CR33938* and *Syp*) (Figure S1I-J), highlighting the comprehensive cell-type representation of our dataset.

### Imp and Syp bind mRNAs encoding regulators of neural development

To obtain functional information about Imp and Syp targets, we performed gene ontology (GO) analysis against the brain transcriptome (Full GO enrichment analysis output available in Supplementary Table 2). Both Imp and Syp bind transcripts encoding proteins with diverse molecular functions, with strong enrichments for DNA/RNA-binding proteins involved in gene expression control at transcriptional and post-transcriptional levels, and also cytoskeleton regulators (Figure 2A). For biological processes, many neurogenesis terms were highly enriched for both Imp and Syp targets, covering broad aspects of neural development from neural stem cell differentiation to axon/synapse maturations (Figure 2B). Loss of Imp results in smaller NBs and under-proliferation of NB lineages, while *syp* knockdown results in the opposite overgrowth and larger brain phenotypes (Figure 2C) (Hailstone et al., 2020; Pahl et al., 2019; Samuels et al., 2020b). We hypothesised that Imp and Syp give rise to these phenotypes via their downstream targets. Therefore, we intersected our dataset with a genome-wide RNAi survey for NB self-renewal and proliferation phenotypes (Neumuller et al., 2011). We found 20% of the RNAi screen genes (127 out of 620) were Imp or Syp targets involved in either NB size, NB number, lineage length, or proliferation defects (Figure 2D). No particular phenotype class was enriched against the screen dataset distribution (Figure S2A). Overall, this result illustrates the pervasive role of Imp and Syp across NB lineage maintenance and differentiation.

**Figure 2.**
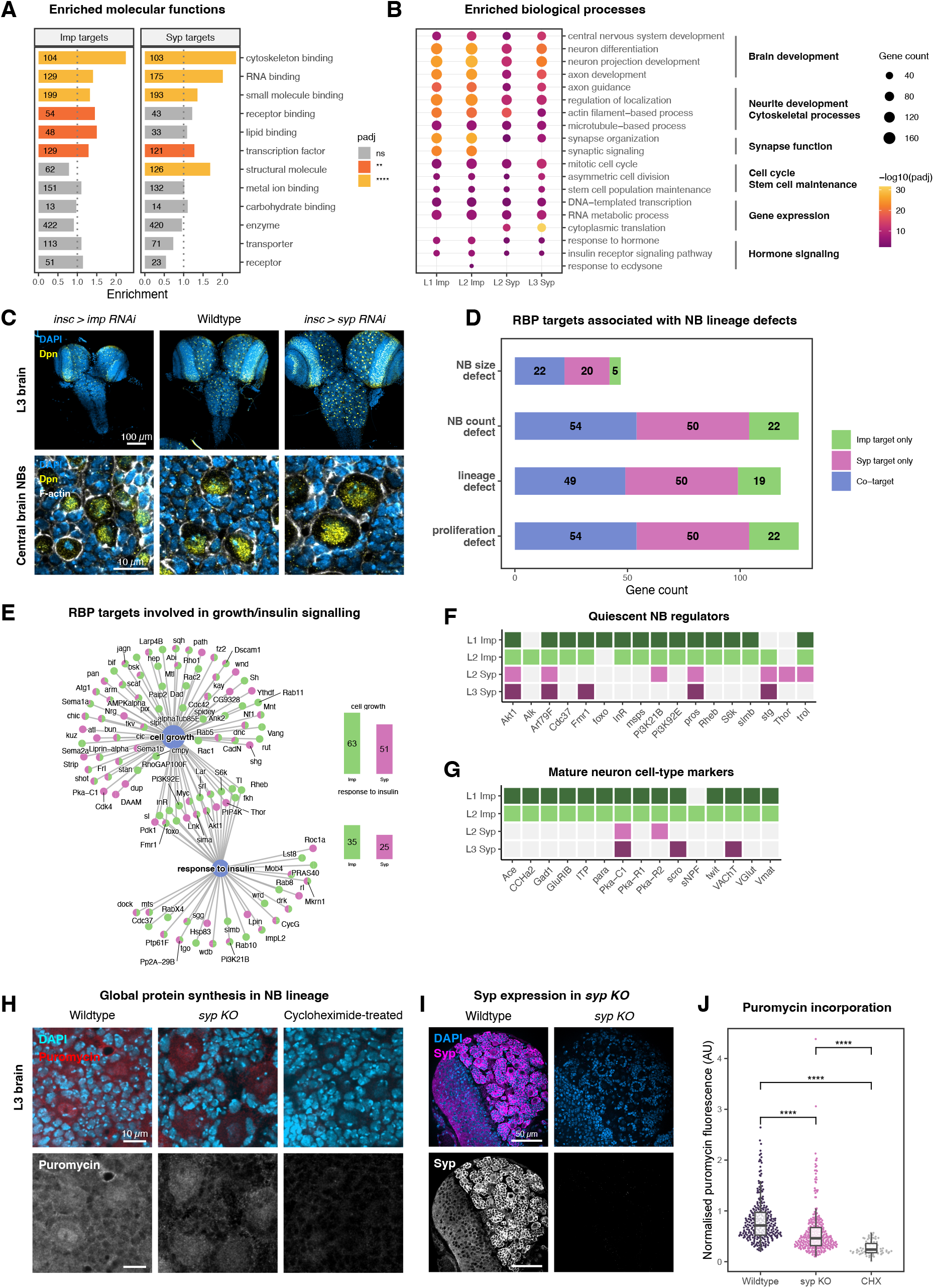
The analysis of Imp and Syp target functions in the neuroblast lineage. **(A)** Enrichments of gene ontology (GO) slim molecular function categories for Imp and Syp targets. The GO categories were taken from the Flybase functional ribbon annotations. Genes from modENCODE larval brain RNA-seq with >10 TPM were used as background. The significance of enrichment was adjusted using Bonferroni’s method. **p<0.01, ****p<0.0001. **(B)** Dotplot showing the top enriched GO:Biological Process terms of Imp and Syp iCLIP targets. Semantically similar GO terms were simplified using the ‘binary cut’ algorithm implemented in the *GOSemSim* R package. Symbols are not shown for non-significant (adjusted p>0.05) terms. **(C)** Comparison of L3 brain and NB size between Wildtype versus *imp* or *syp* knockdown brains (using RNA interference (RNAi)). *insc-GAL4* driver is active in NBs and immediate progenies. anti-Deadpan (Dpn) was used as an NB marker and fluorescently labelled phalloidin was used to mark NB cell boundaries. **(D)** The number of Imp and Syp targets that regulates NB size, NB number, NB lineage length, or neural progenitor proliferation, as identified from a genome-wide RNAi screen (Neumuller et al., 2011). **(E)** Genes annotated with ‘cell growth’ and ‘response to insulin’ terms, grouped by their Imp/Syp target status. Imp and Syp targets are shown in green and magenta, respectively. Bar plots show the number of Imp or Syp targets associated with each term. **(F)**Imp and Syp targets that encode regulators of NB quiescence. **(G)**Imp and Syp targets that encode cell type-specific markers of mature neurons. **(H)**Assessment of puromycin incorporation in Wildtype (*OrR*), *syp knockout (KO)* (*syp* null allele / *syp* deficiency) and the Wildtype brain treated with cycloheximide at the L3 stage. The incorporated puromycin was visualised using anti-puromycin immunofluorescence (see Methods). **(I)** Depletion of Syp protein in the L3 *syp KO* brain compared to the Wildtype. **(J)** Quantification of puromycin incorporation in L3 central brain type 1 NBs from (H). (n=3). ****p<0.0001.

Although Imp and Syp targets were generally enriched for similar biological processes (Figure 2B), Imp targets were more strongly enriched for ‘cell growth’ and ‘response to insulin’ terms (Figure 2E). Cellular growth and insulin signalling are pivotal processes involved in reactivation of quiescent cells (Chell and Brand, 2010; Homem and Knoblich, 2012), and Imp has been shown to influence the timing of NB reactivation in early larval stages (Munroe et al., 2022). Indeed, the majority of the NB quiescence regulators are identified as Imp targets (Figure 2F), suggesting Imp may act on NB reactivation through these transcripts. Furthermore, we found an enrichment of mature neuron cell-type markers (Dillon et al., 2022; Li et al., 2017) (Figure 2G) and ‘synaptic signalling’ GO terms within Imp targets (Figure 2B), reflecting the spatial expression pattern of Imp in functional neurons of embryonic origin. On the other hand, we identified a large group of transcripts encoding ribosomal proteins that bind specifically to Syp (Figure S2B), suggesting its role in regulating ribosome processing and translation. To test whether Syp affects global translation levels, we performed an *ex vivo* puromycin incorporation assay that is sensitive to cycloheximide (CHX) treatment (Figure 2H). Compared to wildtype L3 brains, *syp KO* (*syp^e00286^/Df-BSC124*) brains showed reduced puromycin incorporation, indicating down-regulated protein synthesis in absence of Syp (Figure 2I-J). Taken together, our analysis reveals broad functions of Imp and Syp targets in the nervous system, as well as key differences of Imp and Syp-specific target transcripts.

### Imp and Syp dynamically occupy transcripts encoding temporal factors over time

The post-embryonic neural development programme is heavily influenced by the relative levels of Imp and Syp (Doe, 2017; Islam and Erclik, 2022). Hence, we hypothesised that RNAs involved in temporal fate patterning and NB behaviour may dynamically interact with Imp and Syp over time. To investigate this, we performed a *k*-means clustering based on iCLIP scores of each target that were normalised by their expression levels (Full analysis output available in Supplementary Table 3). Here, we identified six groups of genes displaying distinct binding profiles to Imp and Syp across developmental time points (Figure 3A). As expected, Imp targets were enriched in Clusters I and II (‘Higher Imp binding’) and Syp targets in Clusters III and IV (‘Higher Syp binding’) (Figure 3B).

**Figure 3.**
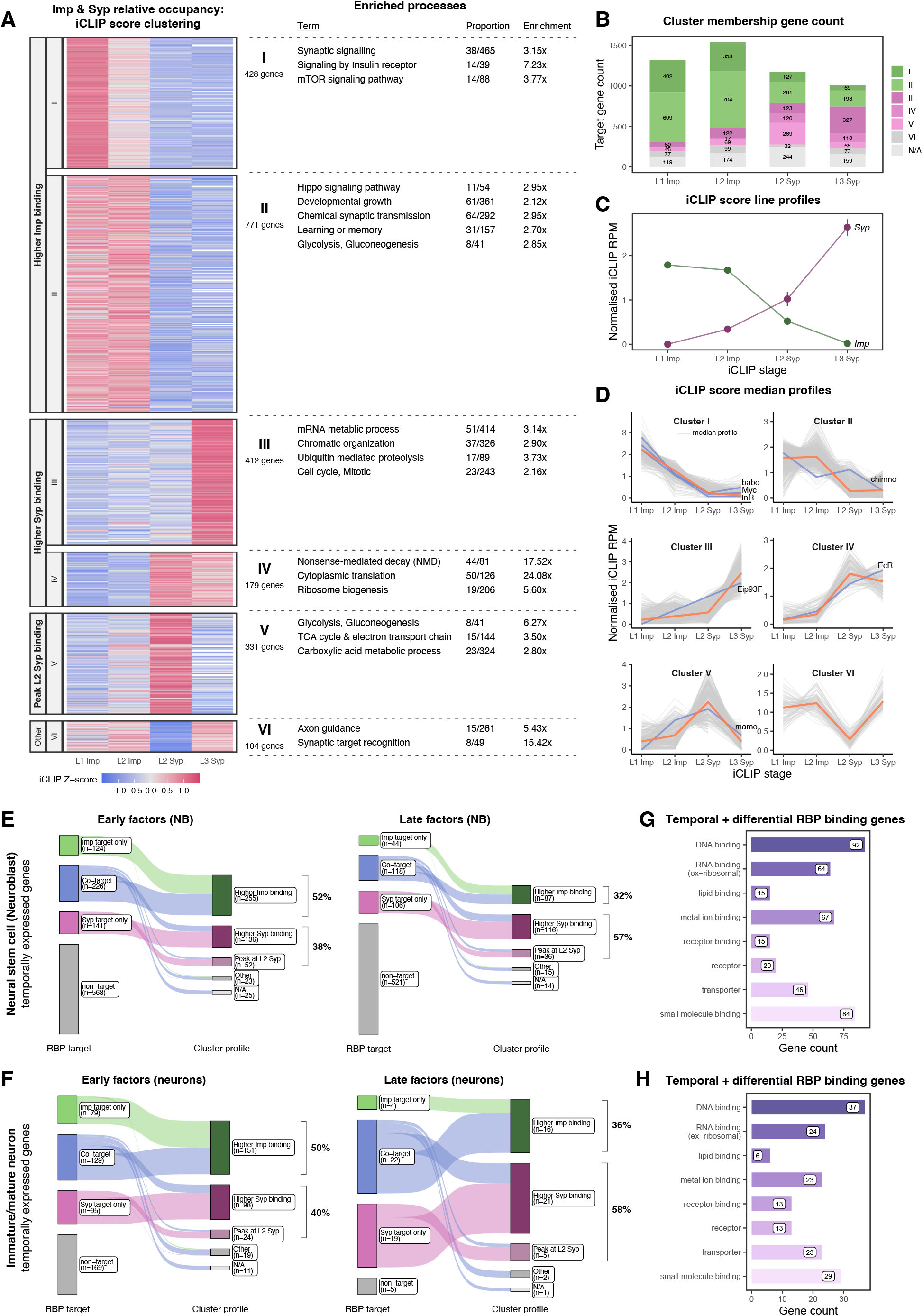
Temporally expressed transcripts interact dynamically with Imp and Syp. **(A)** Heatmap showing *k*-means clustering of Imp and Syp relative occupancy (iCLIP score) at different developmental stages. Relative occupancy for each gene was calculated by normalising iCLIP crosslink counts by the transcript abundance. For each Cluster, enriched GO, KEGG and REACTOME terms are shown. Based on the similarity of the trend, Clusters I and II were grouped into ‘High Imp binding’ group, and Clusters III and IV were designated ‘High Syp binding’. **(B)** Distribution of individual iCLIP library targets by their *k*-means Cluster membership. ‘High Imp binding’ and ‘High Syp binding’ groups are indicated with shades of green and magenta, respectively. **(C)** Line profiles of Imp and Syp relative occupancy to their own transcripts, showing opposing gradients of the interaction. **(D)** Grouped line plots showing iCLIP score profiles of each *k*-means Cluster. Red lines indicate median profiles. The profiles of early temporal factors (*babo*, *Myc, InR,* and *chinmo*) are shown in the ‘High Imp binding’ group, while late temporal factors (*Eip93F* and *EcR*) are shown in the ‘High Syp binding’ group. *mamo*, which specifies the middle mushroom body neuronal fate, is found in Cluster V that shows strong interaction with Syp specifically at the L2 stage. **(E)** Sankey plot showing the intersection between temporally expressed factors in neuroblasts (early and late) and transcripts that are dynamically occupied by Imp and Syp. **(F)** Sankey plot showing the intersection between temporally expressed factors in immature/mature neurons (early and late) and transcripts that are dynamically occupied by Imp and Syp. Temporally expressed transcripts were taken from isolated NB RNA-seq or staged larval brain scRNA-seq (Dillon et al., 2022; Liu et al., 2015; Ren et al., 2017). (**G-H**) Enriched molecular function of temporally expressed genes that dynamically interact with Imp and Syp in (**G**) NB and (**H**) immature/mature neurons. FlyBase GOSlim are shown for high-level classification.

However, some Imp targets were also classified as ‘High Syp binding’ and vice versa, indicating that Imp and Syp can share RNA targets with widely varying occupancy levels (Figure 3B). Each cluster was associated with distinct biological processes as shown by GO, Reactome, and KEGG enrichment analyses (Figure 3A). For example, growth (Hippo, mTOR pathways) and signalling-related transcripts had higher Imp occupancy, while RNA metabolising genes showed stronger interaction with Syp. Interestingly, we found energy metabolism terms were over-represented in ‘Higher Imp binding’ and in Cluster V genes (Figure 3A). Cluster V transcripts strongly interact with Syp specifically at the L2 stage (‘Peak L2 Syp binding’), which represents a key Imp-to-Syp transition period. We hypothesised genes in the ‘Peak L2 Syp binding’ group might be involved in establishing the early-to-late fate transition. Indeed, metabolic switch and oxidative phosphorylation have been shown to impact NB temporal patterning and neuronal specifications (Allen et al., 2020; Davie et al., 2018; van den Ameele and Brand, 2019). Many glycolytic enzymes and components of the TCA cycle showed dynamic interaction with Imp or specifically with Syp at the L2 stage (Figure S3A-B), suggesting the role of Imp and Syp in metabolic reprogramming. In contrast, genes in Cluster VI, with reduced L2 Syp interactions, were enriched for axonogenesis terms (Figure 3A), which reflects the lack of Syp expression in this subcompartment at the L2 stage (Figure 1A).

Next, we investigated whether our dynamic binding analysis could reveal regulators of temporal fates. We found similar opposing gradients of relative RBP occupancy with *imp* and *syp* transcripts reminiscent of their temporal mRNA expressions (Figure 3C) (Liu et al., 2015). Furthermore, known early temporal factors (e.g. *babo*, *Myc*, *chinmo*) were identifiable in ‘Higher Imp binding’ group while late factors (e.g. *Eip93F*, *EcR*) were found in ‘Higher Syp binding groups’ (Figure 3D). To expand our exploration, we analysed dynamic temporal interactors of Imp and Syp that are differentially expressed in early versus late stages of brain development (Corrales et al., 2022; Dillon et al., 2022; Liu et al., 2015). In NBs, 491 early and 268 late factors were targeted by Imp or Syp (Figure 3E), and notably, 90% of these transcripts exhibited dynamic RBP binding profiles. We observed a similar trend for temporally expressed genes in neurons (Figure 3F), and for both cell types, a greater proportion of early factors were classified as ‘Higher Imp binding’ while most of late factors fell under the ‘Higher Syp binding’ group. Inspection of these temporal genes revealed large numbers of DNA and RNA-binding genes in these groups (Figure 3G-H and Figure S3C-D), suggesting that these genes may act directly downstream of Imp and Syp to influence gene expression and subsequent temporal fates.

Neuroblast-derived brain tumours also exhibit a cellular patterning akin to the developmental patterning. For example, *pros*-RNAi induced tumours progress to generate a hierarchy of fast-dividing Imp^+^/Chinmo^+^ cells and slow-dividing Syp^+^/Eip93F^+^ cells (Genovese et al., 2019). Furthermore, both Imp and Syp have been shown to affect proliferation of *brat*-RNAi mediated tumours (Landskron et al., 2018). To contextualise our results, we compared Imp^+^/Chinmo^+^ versus Syp^+^/Eip93F^+^ tumour marker genes on their Imp/Syp relative occupancy group memberships. First, we discovered that a significant number of tumour cell markers turned out to be Imp and Syp targets (Figure S3E). Here, Imp^+^/Chinmo^+^ marker genes were enriched for the ‘Higher Imp binding’ group, while a greater proportion of Syp^+^/Eip93F^+^ markers consisted of ‘Higher Syp binding’ group members (Figure S3E). Notably, the majority of the dynamically interacting tumour markers were already identified in the developing NB/neurons (Figure S3F-G). Considering that metabolic reprogramming also affects tumour immortalisation (Bonnay et al., 2020; Genovese et al., 2019), this result suggests that Imp, Syp, and their targets could be redeployed when establishing tumour differentiation trajectories.

### Imp and Syp post-transcriptionally regulate temporally expressed factors

Next, we examined whether Imp and Syp can regulate their downstream targets. We analysed changes in the target transcript abundance in *syp KO* L3 brain RNA-seq (Samuels et al., 2020a), a system in which the loss of Syp results in protracted expression of *imp* (Figure S4A). We found that ‘Higher Syp binding’ transcripts were generally down-regulated in *syp KO* brains, whereas ‘Higher Imp binding’ genes were up-regulated (Figure 4A). Importantly, classification just by Imp or Syp target status showed relatively uniform distributions of up and down-regulated genes (Figure S4B), indicating that RBP relative occupancy is a better predictor of regulatory trends than the target status alone.

**Figure 4.**
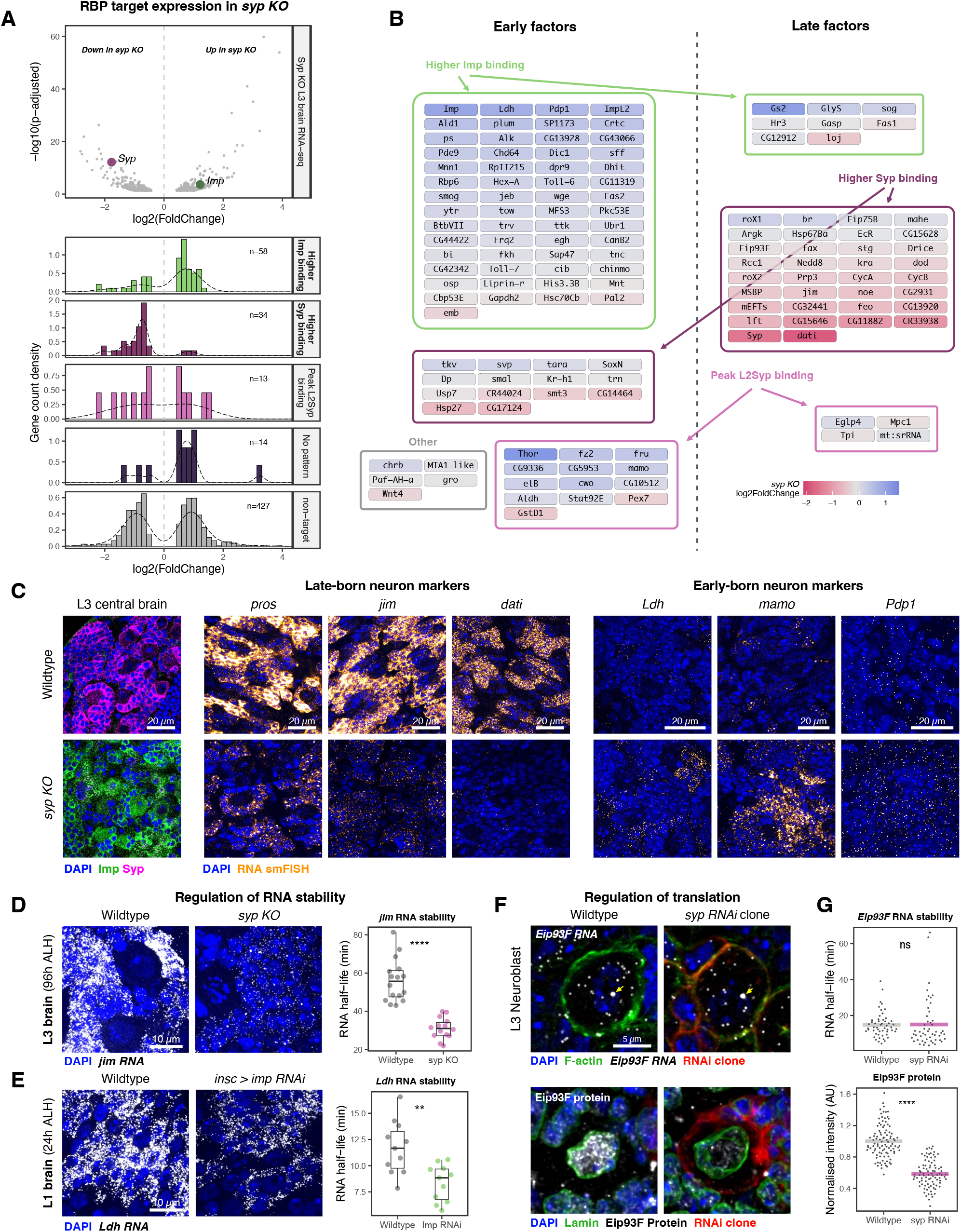
Imp and Syp post-transcriptionally regulate downstream targets. **(A)** RNAs that interact dynamically with Imp and Syp are differentially expressed in L3 *syp KO* brains. In *syp KO* brains, *syp* is downregulated (magenta symbol on the volcano plot) and *imp* is upregulated (green symbol). The histograms show the distribution of log2 fold change in transcript abundance for each RBP-occupancy group. The *syp KO* brain RNA-seq data was taken from (Samuels et al., 2020a). **(B)** A network of early and late temporal factors that dynamically interact with Imp and Syp. The fill colour represents log2 fold change in transcript expression in the *syp KO* brain. Temporal factors were taken from staged larval brain scRNA-seq (Dillon et al., 2022). Transcripts that do not significantly change their expression are also shown. **(C)** Changes in mRNA expression of late-born (*pros, jim, dati*) and early-born (*Ldh, mamo, Pdp1*) immature neuronal markers in Wildtype and *syp KO* L3 brains. Expression of Imp and Syp proteins in Wildtype and *syp KO* brains are shown in the left panel. Type I NB lineages in the central brain regions are shown. Transcript abundance was visualised using smFISH against exon sequences of the mRNAs. **(D)** Visualisation and quantification of *jim* mRNA stability in absence of *syp*. RNA half-lives were measured by relating nascent and mature RNA levels in the L3 (96h ALH) central brain regions (see Methods). (n=3). ****p<0.0001. **(E)** Visualisation and quantification of *Ldh* mRNA stability in *imp RNAi* temporal knockdown system. RNA half-lives were measured in L1 (24h ALH) central brain regions (see Methods). (n=2). **p<0.01. **(F)** Assessment of *Eip93F* RNA and protein levels upon temporal removal of *syp* in central brain Type I NBs. Yellow arrows indicate nuclear transcription sites of *Eip93F*. Temporal *flip-out GAL4 > syp RNAi* clone system was used to knockdown *syp* for 48 hours between the 48-96h ALH period. **(G)** Quantification of *Eip93F* RNA half-lives and nuclear Eip93F protein levels in central brain type 1 NBs after the temporal knockdown of *syp*. (n=3). ****p<0.0001.

The correlation between transcript abundance and RBP relative occupancy suggests a regulatory network where Imp and Syp control target RNA stability to promote early or late temporal fates (Figure 4B). Indeed, our analysis recovered marker transcripts that are indicative of early-born differentiating neurons (*mamo*, *Ldh*, *Pdp1*) and late-born immature neurons (*pros*, *jim*, *dati*), (Figure 4B and Figure S4C) (Dillon et al., 2022; Nguyen et al., 2023). However upstream regulatory roles of Imp and Syp on these transcripts are not fully known. Using smFISH in *syp KO* L3 brains, we found reduced expression of late-born neuronal markers (*pros, jim, dati*) in *syp KO* NB lineages and upregulation of early-fate transcript markers (*Ldh, mamo, Pdp1*) when Imp levels were sustained in absence of Syp (Figure 4C). Importantly, these changes occurred despite active transcription, as evidenced by nuclear transcription sites, suggesting altered transcript stability. We previously demonstrated that Syp can stabilise *pros* mRNA through the extended 3’UTR regulatory sequences (Samuels et al., 2020a). To characterise a further example, we calculated the RNA half-life of a late-born neuron marker *jim* in *syp KO* L3 brains. We calculated *jim* RNA half-life from smFISH images by relating single molecule counts of nascent and mature mRNAs (Figure S4D-E) (Halpern and Itzkovitz, 2016; Mueller et al., 2013), which yielded a value comparable to an orthogonal metabolic labelling method (Thompson et al., 2023). Here, we found significant destabilisation of *jim* in *syp KO* brains (Figure 4D and S4F), suggesting that Syp is crucial for the correct post-transcriptional regulation of *jim*. Similarly, we tested the role of Imp in regulating an early-born neuron marker *Ldh* in the L1 brain (24h ALH). We used a temporal *imp* knockdown system using *insc-*GAL4 and *tub-*GAL80ts (Figure S4G) because constitutive knockdown of *imp* results in NB reactivation defects (Munroe et al., 2022). In this system, loss of *imp* resulted in the destabilisation of *Ldh* in the NB lineage (Figure 4E and S4H). These results suggest that Imp and Syp can influence specification of early or late neuronal fates through regulating RNA stability of their downstream targets.

Imp and Syp can also affect mRNA translation (Liu et al., 2015; Titlow et al., 2020), and this mode of control is not detectable from mutant transcriptomics studies. To characterise an example of regulated translation contributing to temporal patterning, we assessed the gene expression level of *Eip93F*, a temporal TF that interacts genetically with *syp* for late-stage NB specification (Pahl et al., 2019; Syed et al., 2017). *Eip93F* mRNA interacts strongly with Syp (Figure S4I), however its abundance does not significantly change in *syp KO* brains, which suggests invariant RNA stability. Because *Eip93F* transcription in NBs is sensitive to Syp during the Imp-to-Syp transition stage (Figure S4J), we assessed mRNA half-life and protein level of Eip93F in the temporal *syp* knockdown system (Figure S4K-M). We found comparable RNA stability of *Eip93F* between Wildtype and *syp* RNAi clone NBs, but Eip93F protein was significantly down-regulated in the absence of Syp (Figure 4F-G). This indicates Syp is required for the correct protein synthesis output of *Eip93F*. Taken together, our results suggest that Imp and Syp are key post-transcriptional regulatory effectors in fate patterning. We propose many more transcripts that dynamically interact with these RBPs may play a role in specifying coarse or fine temporal windows.

### Competitive binding interplay between Imp vs Syp is rare

Distinct signatures of Imp and Syp occupancy on target transcripts indicate that their molecular interplay plays a crucial regulatory role. Therefore, we next investigated whether iCLIP peak distributions could reveal regulatory modalities of the interplay between Imp and Syp. First, we considered a scenario where Imp and Syp directly compete for the same binding sites. To address whether Imp and Syp can recognise similar RNA sequences, we performed a motif enrichment analysis by comparing iCLIP peaks versus background crosslinks (Kuret et al., 2022). As expected, given the 3’UTR preference, Syp binding sites were highly enriched in AU-containing motifs, potentially part of the regulatory AU-rich elements (AREs) (Figure 5A). Imp, at both stages however, predominantly bound to CA-rich motifs, though a small number of AU-rich motifs scored above the background (Figure S5A). Consistently, PCA of *k*-mer enrichment scores revealed a clear distinction between Imp and Syp binding motifs across mRNA and ncRNA features (Figure 5B and Figure S5A-B). We also examined individual motif enrichments at the L2 stage where both RBPs co-express and spatially coincide, yet it revealed a poor correlation (R^2^ = 0.019) between Imp and Syp binding sites (Figure 5C), indicating a lack of consensus sequence motifs that are commonly recognised.

**Figure 5.**
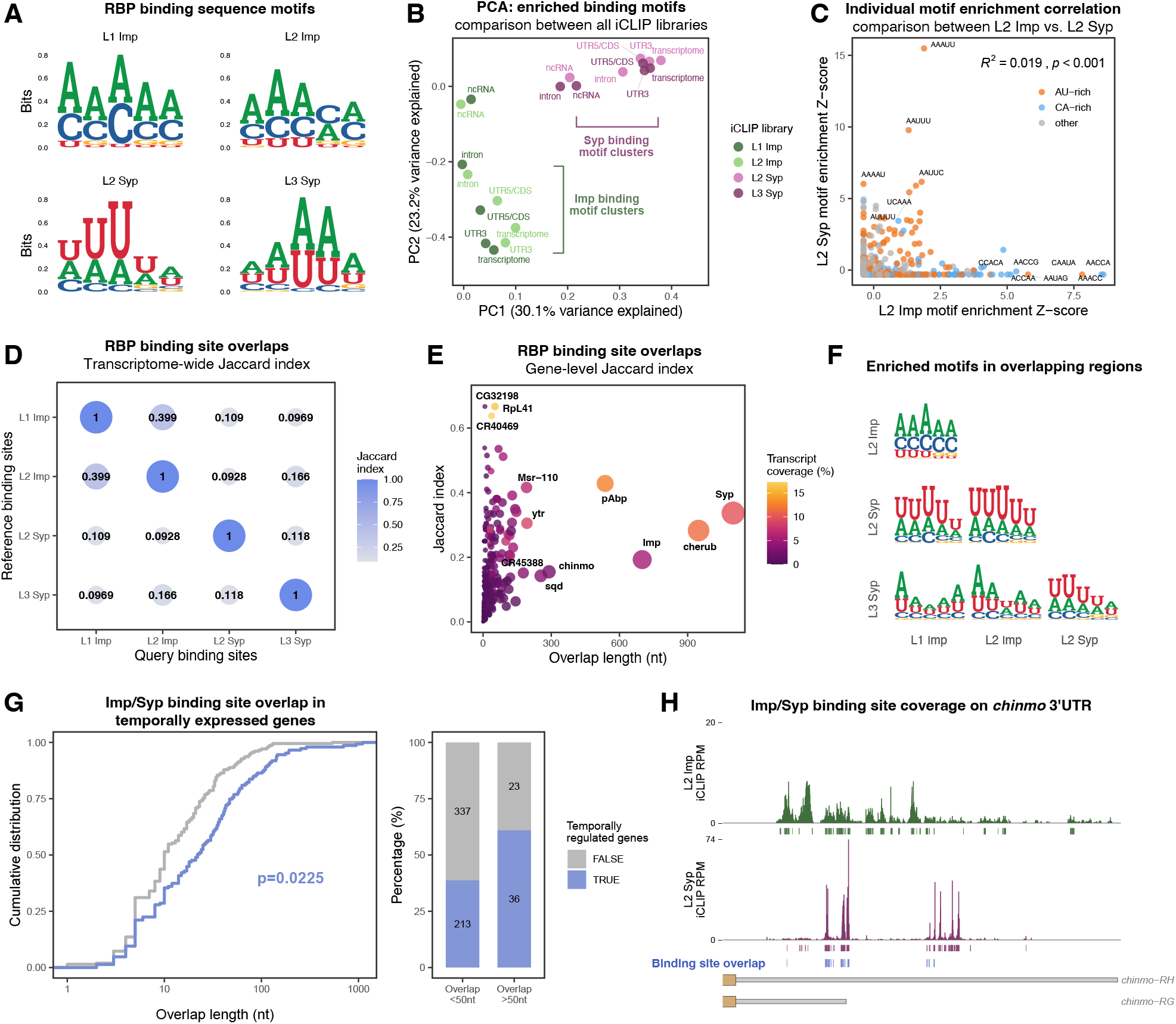
Imp and Syp binding sites poorly overlap, except in temporally expressed transcripts. **(A)** Enriched RNA sequence motifs of Imp and Syp binding sites. Non-significant crosslinks (FDR > 0.01) were used as backgrounds using PEKA (see Methods). The consensus motifs obtained across all transcriptomic regions are shown. **(B)** Principal component analysis (PCA) of enrichment scores of all sequence motifs (5-8 *k*-mers) from Imp and Syp iCLIP libraries. Enrichment scores are further divided into transcriptomic regions. Note the separation of Imp and Syp libraries, while L1/L2 Imp and L2/L3 Syp motifs cluster closely with each other. **(C)** Correlation of individual sequence motif enrichment scores between L2 Imp and L2 Syp iCLIP libraries. AU-rich or CA-rich motifs are coloured separately, and R^2^ correlation value is shown. **(D)** Transcriptome-wide overlap between Imp and Syp iCLIP binding sites. The degree of overlap was assessed using the Jaccard index method. **(E)** Gene-level overlap between L2 Imp and L2 Syp iCLIP binding sites. The size of the symbol represents the absolute nucleotide-length of the overlap. **(F)** Enriched RNA binding sequence motifs specifically in the overlapping binding sites between Imp and Syp iCLIP libraries. **(G)** Cumulative distribution comparing L2 Imp and L2 Syp iCLIP binding site overlap between temporally regulated and non-temporally regulated genes (*left*). Bar plot showing the group of temporally expressed genes and whether Imp & Syp binding sites overlap more than 50nt on their mRNA. Temporally regulated genes were taken from staged NB and brain scRNA-seq datasets (Dillon et al., 2022; Liu et al., 2015; Ren et al., 2017). **(H)** Imp and Syp iCLIP coverages on the *chinmo* 3’UTR region. Overlapping binding sites are highlighted in blue.

To explore this further, we investigated the physical relationship between Imp and Syp footprints on their co-targets. At a global scale, we found a statistically significant proximity between Imp and Syp iCLIP peaks (Figure S5C). However, direct overlap measurements yielded very small Jaccard indices, suggesting an unlikely coincidence of overlapping binding events (Figure 5D). We repeated this analysis at the individual gene level and also found a generally weak overlap across most co-targets (Figure 5E). However, we identified an outlier group of transcripts (5%) with notable degrees of overlap, many of which are known to be regulated temporally, such as *chinmo*, *lncRNA:cherub*, *sqd* as well as *imp* and *syp* themselves (Figure 5E). Within these regions, AU-rich sequence motifs were more enriched over CA-rich motifs (Figure 5F). We hypothesised temporal genes under regulatory influence of Imp and Syp could experience direct competition between the two RBPs. To test this, we compared coverage of overlapping binding sites between temporal and non-temporal genes identified from scRNA-seq or mechanically dissociated NB transcriptomes (Corrales et al., 2022; Dillon et al., 2022; Liu et al., 2015; Ren et al., 2017). We found that temporal genes harbour longer regions of Imp and Syp footprint overlap than non-temporal genes (Figure 5G). Of note, as illustrated with *chinmo*, the absolute extent of overlaps was still modest in most genes, where only 17% of the RBP coverages were coincidental (Figure 5H). However *syp* was a notable outlier with 35% coincidental footprints (Figure 5E). We propose that a rare subset of temporally-regulated transcripts may experience direct competition between Imp and Syp for the same binding sites, though this competition is unlikely to be a globally deployed regulatory mode.

### Imp and Syp binding sites show co-evolutionary and combinatorial binding signatures

To further characterise the functional importance of Imp and Syp binding to their target transcripts, we examined whether their binding sites are evolutionarily conserved. We calculated phastCons sequence conservation score (across 27 *Drosophila* species) (Siepel et al., 2005) of iCLIP peaks and found that Imp and Syp binding regions were significantly more conserved than the brain transcriptome (Figure 6A and Figure S6A). To rule out targeting bias towards conserved genes, we calculated average phyloP sequence conservation scores for each transcript feature in each gene and compared these with shuffled RBP binding sites per queried region (Pollard et al., 2010). We found Imp and Syp binding sites were highly conserved across multiple transcript features, except for CDS in Imp and ncRNA in Syp (Figure 6B). Similar conservation trends were also observed when expanding the analysis to include 124 insect species (Figure S6A-D), indicating a functional importance of Imp and Syp binding in maintaining evolutionary fitness.

**Figure 6.**
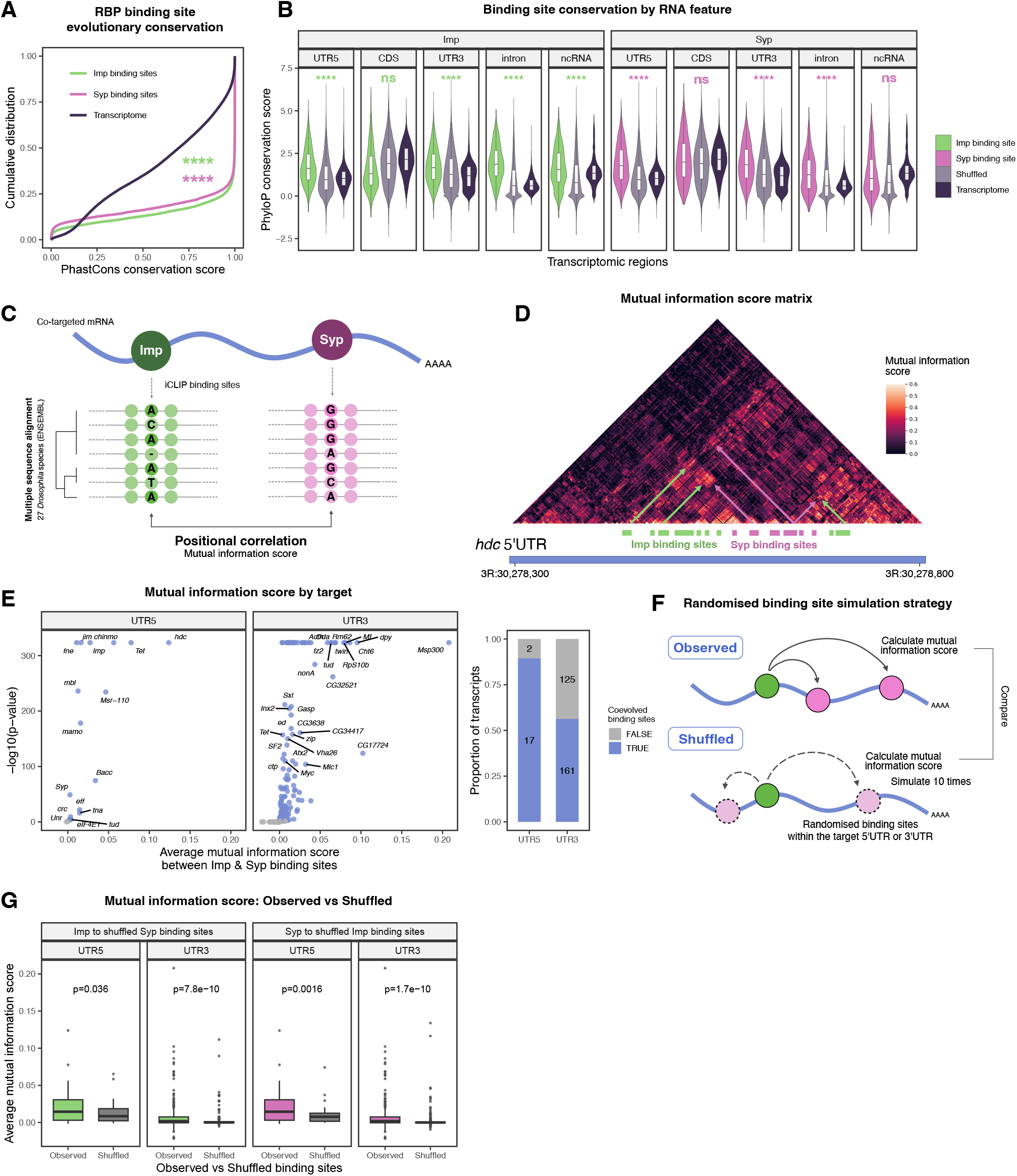
Imp and Syp binding sites display evolutionary linkage. **(A)** Comparison of PhastCons RNA sequence conservation score of Imp and Syp binding sites versus the larval brain transcriptome average. 27-way *Drosophila* PhastCons track was taken from the UCSC genome browser. ****p<0.0001. **(B)** Analysis of PhyloP RNA sequence conservation score of Imp and Syp binding sites per gene and per transcript feature. The same scores calculated for the brain transcriptome and the shuffled iCLIP binding sites were used as controls. PhyloP was used instead of PhastCons due to its independence scoring between short-range nucleotide distances. For shuffled binding sites, the simulation was iterated 10 times per query. ****p<0.0001. **(C)** A strategy to calculate mutual information (co-evolution) score between Imp and Syp binding sites. To limit the analysis to confident binding sites, iCLIP peaks present in both L1/L2 for Imp and L2/L3 for Syp were used. **(D)** A correlation matrix of mutual information score between every nucleotide pair in the *headcase* (*hdc*) 5’UTR. Local clusters of high mutual information score coincide with Imp and Syp binding site pairs. **(E)** Average mutual information (co-evolution) scores of Imp and Syp binding sites per gene. For each target gene, the evolutionary linkage was assessed by comparing average mutual information score between Imp and Syp binding site pairs versus their column average. Adjusted p-value < 0.01 was considered significant. Due to computational memory constraints, mutual information calculations were subdivided between 5’UTR and 3’UTR. **(F)** Control simulation experiment for mutual information score calculation. The schematics illustrate a strategy to compare the distribution of average mutual information scores in *observed* (empirical iCLIP binding sites) versus randomised binding sites (*shuffled*). The simulation was performed 10 times per gene per UTR region and in both Imp-to-Syp and Syp-to-Imp directions. **(G)** Comparison of average mutual information scores in *observed* and *shuffled* settings, as described in (F).

Expanding on the RBP footprint conservation, we next explored whether Imp and Syp binding sites jointly mutated to maintain genetic interactions. Given the limited overlap between Imp and Syp binding sites, we hypothesised that their combinatorial mode of molecular interplay could be detected through evolutionary couplings. (Weinreb et al., 2016). To investigate this, we performed mutual information analysis using a phylogenetic inference across 27 *Drosophila* species to identify pairs of co-evolving nucleotides within target mRNA sequences (Figure 6C) (Alfonso-Gonzalez et al., 2023b; Brandman et al., 2012; Martin et al., 2005). We calculated mutual information scores across 5’UTR or 3’UTR of the shared RNA targets of Imp and Syp at the L2 stage where the RBPs spatially coexpress (Full analysis output available in Supplementary Table 4). We found local clusters of high-scoring nucleotide pairs that frequently coincided with Imp and Syp binding site pairs (illustrated with the *headcase* (*hdc*) 5’UTR score matrix, Figure 6D), which is suggestive of co-evolutionary linkage between their binding sites. Globally, we observed significantly higher average mutual information scores between Imp and Syp footprint pairs compared to individual row averages in 89% and 56% of the 5’UTR and 3’UTR targets, respectively (Figure 6E). To reinforce our finding, we performed a control simulation experiment. We calculated co-evolution scores upon randomised shuffling of the partner RBP binding sites and simulated this ten times for both Imp-to-Syp and Syp-to-Imp directions (Figure 6F). In this scenario, the significantly lower average mutual information scores in the *shuffled* simulations compared to the experimentally *observed* values further supported the pairwise evolutionary interaction between Imp and Syp footprints (Figure 6G). Overall, our results show Imp and Syp binding sites are highly conserved and likely jointly mutated to maintain their regulatory link. Therefore, we propose that combinatorial interplay, rather than competitive binding, between Imp and Syp could be more widespread in producing correct regulatory outcomes for their downstream co-targets.

## Discussion

Here, we identify transcriptome-wide targets of Imp and Syp, a pair of RBPs that exhibit the most dynamic temporal expression gradient during the post-embryonic brain development. RBPs control gene expression by regulating the RNA metabolism of their targets, making the elucidation of their RNA interactomes crucial for understanding their core functionalities. Temporal patterning in larval and pupal stages requires a combination of transcription factors and the opposing gradients of Imp and Syp (Guan et al., 2022; Liu et al., 2015; Syed et al., 2017; C. P. Yang et al., 2017). Remarkably, more than 60% of VNC neurons are born during this period, with their initial patterning at this stage, pre-configuring adult neuronal identity (Nguyen et al., 2024). Much work has been done to understand the chromatin-level control of temporal specification. However, discordant transcription and translation activity of terminal differentiation genes (e.g. neurotransmitters) suggests RNA regulation can also play a substantial role (Marques et al., 2023). Our high-resolution study of RBP target landscapes and their RNA footprints contributes to understanding the connections between concurrent transcriptional and post-transcriptional mechanisms shaping the nervous system.

Opposing Imp and Syp temporal gradients have been described to pattern multiple post-embryonic lineages in the central brain, VNC and optic lobe (Arain et al., 2022; Liu et al., 2015; Ren et al., 2017; Syed et al., 2017). While the slopes of Imp and Syp expression vary considerably between lineages, it is likely that the RBP gradient assigns coarse fates in these lineages, with temporal TF cascades downstream of Imp and Syp further diversifying neuronal subtypes (Islam and Erclik, 2022). Our temporal RBP occupancy analysis highlights a group of transcripts, enriched for TFs and RBPs, that differentially interact with Imp and Syp in a temporal manner. These transcripts are good candidates for defining fine temporal sub-windows. In other respects, unique modes of division deployed in Type II INP and optic lobe lineages indicate that spatial patterning mechanisms act concurrently with temporal gradients in order to diversify neural fates (Bayraktar and Doe, 2013; Konstantinides et al., 2022). While this study aimed to identify brain-wide targets of Imp and Syp, identifying cell-type specific RNA interactomes may facilitate future studies on the cooperation between multiple patterning mechanisms. For example, HyperTRIBE technique, which utilises UAS-RBP::ADAR fusion constructs, will be useful to decipher lineage or cell-type specific RBP targets (Rahman et al., 2018).

Our dataset captures Imp and Syp targets involved in diverse biological and molecular functions, including key regulators of NSC growth and quiescence, energy metabolism, intracellular signalling, hormone response, and tumourigenic potential. This indicates the versatility of Imp and Syp gradients in mediating intrinsic NB lineage functions as well as responding to extrinsic signals (Branham et al., 2024; Islam and Erclik, 2022; Rossi and Desplan, 2020; Syed et al., 2017). Notably, both Imp and Syp bind to many transcripts encoding cytoskeleton and axonogenesis regulators (Figure 2A). Despite their known roles in synaptic transmission and cytoskeletal remodelling, the developmental context of Imp and Syp in circuit connectivity is not well understood (Halstead et al., 2014; Hansen et al., 2015; Titlow et al., 2020; Vijayakumar et al., 2019). In the central brain Type I lineage, early-born neurons form extensive neurite projections while later-born neurons display decreasing morphological complexity (Lee et al., 2020). Both Imp and Syp proteins localise to nerve bundles and distal tips of immature neurites, therefore it would be interesting to investigate the regulatory function of Imp and Syp within these fine neuronal projections.

Given their predominantly cytoplasmic localisation, Imp and Syp likely regulate the stability, translation and/or localisation of their downstream target transcripts (Di Liegro et al., 2014; McDermott et al., 2014; Munro et al., 2006). This is reflected in the enrichment of iCLIP binding sites in the 5’ and 3’ UTRs, which harbour *cis*-regulatory sequences that may respond to changing Imp and Syp levels (Dillard et al., 2018; Samuels et al., 2020a). A key example is *chinmo*, where Imp promotes translation while Syp represses translation, resulting in a steep gradient of the protein product despite a relatively constant level of its mRNA (Liu et al., 2015; Zhu et al., 2006). The substantial overlap between Imp and Syp RNA interactomes suggests they may influence divergent expression of many more genes in the brain. Therefore, it is important to consider the mechanistic interplay between *trans*-acting factors as a critical layer of regulation (Van Nostrand et al., 2020). For example, regulatory interplay between RBPs can be competitive when they occupy overlapping binding sites, causing steric hindrance (Dassi, 2017). However, our RBP profiling method does not support this as the primary mode of interplay, as Imp and Syp recognise distinct sequence motifs with limited binding site overlaps. Nevertheless, these small overlapping regions might still act as key regulatory hubs for direct competition and regulatory outcomes (Tiedje et al., 2012; Topisirovic et al., 2009). Future mutagenesis studies are needed to determine whether these overlapping binding sites are essential for the transcript’s response to the opposing gradients of Imp and Syp.

Alternatively, RBPs can bind to the same RNA in a combinatorial manner, where differential occupancy can have synergistic or antagonistic effects on target gene expression (Schueler et al., 2014; Van Nostrand et al., 2020). This mode of interplay also includes RNA-dependent interactions, such as binding of one RBP triggering allosteric changes in the RNA structure that facilitates or hinders the binding of another RBP (Iadevaia and Gerber, 2015; Nag et al., 2022). Our binding site coevolution analysis supports the combinatorial interplay between Imp and Syp and the conservation of the Imp-Syp regulatory cassette in many co-targeted mRNAs. In this context, Imp and Syp likely regulate the fate of the bound mRNA by recruiting other effector proteins or forming processive multi-RBP complexes. In mammals, IGF2BP1 associate with HNRNPU, SYNCRIP, YBX1, and DHX9 to control *c-myc* stability, and hnRNPQ has been shown to interact with PABP to regulate cap- and IRES-mediated translations (Degrauwe et al., 2016; Svitkin et al., 2013; Weidensdorfer et al., 2009). Consistent with this idea, our dataset shows over-representation of transcripts encoding RBPs targeted by Imp and Syp. This suggests that Imp and Syp could influence expression of co-interacting partner RBPs to tailor their post-transcriptional regulatory niche. It would be important to elucidate protein interactors of Imp and Syp in the brain and how combinatorial binding of Imp and Syp affects recruitment of these factors. While, in our knowledge, this is the first report of iCLIP in larval tissues, future RBP profiling experiments will be valuable to investigate binding site co-occurrences across RBPs and discover novel co-interacting RBPs.

Mammalian homologs of Imp and Syp are also expressed in the developing brain and play crucial roles in neuro/synaptogenesis (Chen et al., 2012; Mori et al., 2001; Williams et al., 2016). For example, in mice, IMP1 levels rapidly decline in a temporal fashion to regulate maintenance and differentiation of vertebrate NSCs (Nishino et al., 2013). Likewise, *snRNA:7SK*, a high target of fly Syp, has been shown to interact with hnRNP-R to regulate local transcriptome in neurites (Briese et al., 2018). In the current study, we found a surprising degree of evolutionary conservation in RNA target preference and RNA sequence recognition between invertebrate and vertebrate orthologues of Imp and Syp. Morphogenetic gradients and temporal cascade of RBPs and transcription factors are likely conserved in mammalian neurogenesis to specify young and old cellular fates (Caviness et al., 2009; Doe, 2017; Mattar et al., 2015; Telley et al., 2019; Zahr et al., 2018). In the future, it will be interesting to determine if orthologous temporal factors downstream of Imp and Syp play conserved roles in the developing vertebrate CNS.

## Materials and methods

### Resource table

| Reagent type or resource | Designation | Source or reference | Identifiers | Additional information |
| --- | --- | --- | --- | --- |
| Gene (*D. melanogaster*) | *Drosophila melanogaster* genome assembly | Ensembl | BDGP6.28 | Release 99 |
| Genetic reagent (*D. melanogaster*) | *Syp::GFP* | (Titlow et al., 2022) |  |  |
| Genetic reagent (*D. melanogaster*) | *Imp::GFSTF* | BDSC | 60237 |  |
| Genetic reagent (*D. melanogaster*) | *Wildtype (OrR)* | BDSC | 5 |  |
| Genetic reagent (*D. melanogaster*) | *insc-GAL4* | (Betschinger et al., 2006) |  |  |
| Genetic reagent (*D. melanogaster*) | *syp RNAi* | VDRC | 33011 |  |
| Genetic reagent (*D. melanogaster*) | *tub-GAL80ts ; TM2/TM6B* | BDSC | 7108 |  |
| Genetic reagent (*D. melanogaster*) | *imp RNAi* | BDSC | 34977 |  |
| Genetic reagent (*D. melanogaster*) | *Control RNAi (Blank)* | VDRC | 60100 | Used for Wildtype control |
| Genetic reagent (*D. melanogaster*) | *syp*<sup>e00286</sup> | BDSC | 95448 | Syp null allele |
| Genetic reagent (*D. melanogaster*) | *syp*<sup>f03775</sup> | BDSC | 85668 | Syp hypomorphic allele |
| Genetic reagent (*D. melanogaster*) | *Df3R(BSC124)* | BDSC | 9289 | Syp deficiency |
| Genetic reagent (*D. melanogaster*) | *hsFLP ; act>Flipout>GAL4, UAS-GFP* | Gift from A. Baena-Lopez |  |  |
| Antibody | anti-Syp (guinea pig, polyclonal) | (McDermott et al., 2014) |  | 1:500 IF |
| Antibody | anti-Imp (rabbit, polyclonal) | Gift from P. Macdonald |  | 1:500 IF |
| Antibody | anti-Eip93F (guinea pig, polyclonal) | Gift from C. Doe |  | 1:500 IF |
| Antibody | anti-Dpn (rat, monoclonal) | Abcam | ab195173 | 1:200 IF |
| Antibody | anti-GFP (rat, monoclonal) | Chromotek | 3H9 | 1:1000 WB |
| Antibody | anti-FLAG M2 (mouse, monoclonal) | Sigma | F1804 | 1:1000 WB |
| Antibody | anti-Puromycin | Sigma | ZMS1016 | 1:50 IF |
| Antibody | anti-Lamin | DSHB | ADL67.10 | 1:10 IF |
| Commercial assay, kit | GFP-trap magnetic agarose | Chromotek | gtma-20 |  |
| Commercial assay, kit | Anti-FLAG M2 magnetic beads | Sigma | M8823 |  |
| Chemical compound, drug | Alexa Fluor 488 Phalloidin | ThermoFisher | A12379 |  |

### Fly genetics

All fly stocks were raised on standard cornmeal-agar medium at 25°C. For iCLIP experiments, larvae were developmentally synchronised at the larval-hatching stage by collecting newly hatched flies within a 2h window into apple juice plates supplemented with yeast paste. GAL4-positive neuroblast clones were generated using the heat-shock inducible flippase (*hsflp*) element and the GAL4 flip-out cassette under the control of a transcription stop sequence flanked by FRT sites. Appropriately staged larvae were heat-shocked at 37°C for 15-30min in a water bath to express *hsflp*, which stochastically induces GAL4-positive neuroblast clones. The heat-shocked flies were maintained at 29°C until dissection.

### iCLIP library preparation

For whole-larval and larval-head iCLIP, samples were collected and snap-frozen in liquid nitrogen. Frozen biosamples were ground into fine powder in a liquid nitrogen-cooled mortar and pestle and were subject to 254nm UV-C irradiation four times at 150-200mJ/cm2 using a Stratalinker 2400 (Stratagene). For larval brain iCLIP, wandering L3 brains (96h after larval hatching, ALH) were dissected in cold PBS supplemented with 0.2mM 4-2-aminoethyl)benzenesulfonyl fluoride hydrochloride (AEBSF) in batches of 50 brains, and UV-C irradiated four times at 150mJ/cm2 with shaking in-between. Crosslinked samples were lysed in CLIP lysis buffer (Buchbender et al., 2019) supplemented with 0.2-2mM AEBSF (Sigma), 1-2x cOmplete protease inhibitor (Roche), 1mM Ribonucleoside vanadyl complex (RVC, New England Biolabs), and 40U/ml RNAsin plus (Promega). Lysates were sonicated using a Bioruptor (Diagenode) with 10 cycles of 30s on/off periods at low intensity setting. Lysates were cleared by centrifugation and filtration using Co-star 0.22μm spin filters (Sigma), and protein levels were quantified using the Pierce 660 Protein Assay (ThermoFisher). Quantified lysates were normalised to 0.5-1.5mg/ml, snap-frozen and stored at −80°C.

iCLIP sequencing libraries were generated using the iCLIP2 protocol with several modifications from the infrared CLIP (irCLIP) and the enhanced CLIP (eCLIP) protocols (Buchbender et al., 2019; Lee and Ule, 2018; Van Nostrand et al., 2016; Zarnegar et al., 2016). Crosslinked RNAs in the lysates were partially trimmed with 0.15-0.5U/μl of RNAse I (Ambion) for 3min at 37°C. GFP-tagged Syp (L3 brains) was immunoprecipitated using GFP-Trap Magnetic Agarose Beads (Chromotek), and Syp (L2 heads) was immunoprecipitated using guinea-pig anti-Syp (McDermott et al., 2012) coupled to Dynabeads Protein A (Invitrogen). FLAG-tagged Imp was pulled down with Magnetic M2 FLAG Beads (Sigma). Immunoprecipitations and washes were performed as described in the iCLIP2 protocol, and the RNAs were 3’end dephosphorylated by PNK (NEB) and FastAP phosphatases (Thermo). Then, infrared IR800-dye conjugated 3’ adapters were ligated overnight using RNA ligase 1 (NEB) in a high-percentage PEG8000 buffer. Molecular weight distribution of the protein-RNA complexes was visualised after 3-8% Tris-Acetate SDS-PAGE and transferred to nitrocellulose membranes, then appropriate size regions (∼75 kDa above the expected molecular weight of the RBPs) were excised to extract RNA using proteinase K (Roche) in an SDS-containing buffer (Zarnegar et al., 2016). The RNAs were further purified with phenol/chloroform/isoamyl alcohol mix (pH 6.6) followed by clean-up with RNA Clean & Concentrator Kit (Zymo research). The purified RNAs were reverse transcribed using SuperScript IV (Invitrogen), and the cDNA was purified with MyONE Silane beads (Thermofisher). Second DNA adapters containing unique molecular identifiers (UMIs) and experimental barcodes were attached to the 3’ end of the cDNAs overnight using RNA ligase 1, and the reaction clean-up was performed using MyONE Silane beads. The cDNAs now flanked by Illumina adapters at both ends were amplified using a two-step PCR strategy. First, cDNAs were amplified with 6 PCR cycles using short Solexa P3/5 primers to allow beads-based size selection with ProNex Beads Size-selection Chemistry (Promega). In order to avoid over-amplification, the optimal number of cycles for the second amplification was calculated using 1μl of the first PCR product from qPCR Ct values. Both PCR and qPCR were performed using Pfusion High-Fidelity Polymerase Master Mix (NEB) and full-length Solexa P3/5 primers, and Evagreen (Biotium) was added as a reporter dye in qPCR reactions. After the second PCR amplification, libraries were subject to a second round of size selection by 2.4X ProNex Chemistry to deplete excess primers and molecules shorter than 150bp.

Size-matched input (SMInput) libraries were prepared in parallel with immunoprecipitated samples. 10μl of the RNAse I digested lysates were separated via SDS-PAGE alongside the paired immunoprecipitation sample. After transfer to a nitrocellulose membrane, the same molecular weight region was excised, hence size-matched, and RNA was extracted, which represents the background RBP crosslink-ome. Purified RNAs were 3’end dephosphorylated and adapter-ligated in solution followed by MyONE Silane clean-up after each step. Subsequently, the SMInput RNAs were processed alongside the immunoprecipitation sample at the reverse transcription stage.

Normalised and pooled libraries were sequenced using a NextSeq 500 sequencer (Illumina) at the Department of Zoology, University of Oxford. High-output cartridge v2.5 was used for 92 cycles in single-end mode.

### Bioinformatics methods

#### iCLIP computational analysis

Raw FASTQ files from iCLIP were pre-processed as described in (Busch et al., 2019) with minor modifications. The reads were mapped to the Ensembl *D. melanogaster* genome assembly version 99 using *STAR* with end-to-end mapping (Dobin et al., 2013). PCR duplicated cDNAs were filtered based on unique molecular identifier (UMI) sequences and mapping locations using *UMI tools* (https://github.com/CGATOxford/UMI-tools). Crosslink sites were extracted, which were defined as the nucleotide position preceding the cDNA insert (i.e one nucleotide before the reverse transcription truncation events). Significant crosslink sites and RBP binding peaks were called using the *iCount* package (https://icount.readthedocs.io) with a false discovery rate (FDR) threshold of 0.01. The binding peaks were further filtered by comparing iCLIP versus SMInput libraries using *DESeq2* (Love et al., 2014) with log2FoldChange > 1 and padj < 0.05 threshold. Finally, significant and enriched binding sites were normalised to 5nt length using the R script provided in (Busch et al., 2019).

#### RNA-seq analysis

Public RNA-seq datasets were re-analaysed using the Kallisto-DESeq2 method (Bray et al., 2016). Raw FASTQ files were downloaded from Gene Expression Omnibus (GEO) and were filtered for fly ribosomal RNA and low-complexity reads using BBDuk (https://github.com/BioInfoTools/BBMap). Filtered reads were pseudoaligned to the Ensembl *D. melanogaster* transcriptome assembly version 99 using Kallisto. The custom transcriptome index was created using *Kallisto-Bustools* to include flanking intron sequences (25nt) at either ends of exons. Estimated counts and transcripts per million (TPM) for transcript isoforms were collapsed for each gene. Differential expression analysis was performed using the R package *DESeq2*. The normalisation for library size was performed within the DESeq2 analysis and multiple hypothesis testing was done via the Independent Hypothesis Weighting (IHW) method (Ignatiadis et al., 2016) implemented in the R package *IHW*. For differential expression of genes in neuroblast temporal datasets (Liu et al., 2015; Ren et al., 2017), the pseudoaligned counts were processed using maSigPro (Conesa et al., 2006) with default settings.

#### Area under recovery curve Cell (AUCell) measurements

scRNA-seq gene expression matrices and associated embedding information were downloaded from *Scope* (https://scope.aertslab.org/) or from (Dillon et al., 2022). To identify cells with active gene sets (expression of Imp and Syp target transcripts), AUCell was used (Aibar et al., 2017). For each cell from the scRNA-seq dataset, AUCell scores for combined Imp targets (L1 + L2) and Syp targets (L2 + L3) were calculated from non-normalised gene expression matrices. The resulting scores represent the proportion of expressed transcripts in the query gene set and their relative expression value compared to the other transcripts in the cell.

#### Functional annotation and enrichment analysis

Gene ontology (GO), Reactome and Kyoto Encyclopedia of Genes and Genomes (KEGG) pathway enrichment analyses were conducted using the R package *clusterProfiler* (Yu et al., 2012). Terms that are log2FoldChange > 1 and adjusted p-value < 0.05 were considered significantly enriched. For ‘Biological Process’ GO terms, similar terms were grouped using the ‘binary_cut’ algorithm using the R package *GOSemSim* (Yu, 2020). For ‘Molecular Function’ GO terms, genes were grouped into GOSlim terms represented in FlyBase gene function ribbon.

#### RBP target comparison between species

Human IMP1-3 eCLIP data from pluripotent stem cells were filtered for crosslink foldchange > 2 to identify IMP targets (Conway et al., 2016). Rat SYNCRIP iCLIP targets from primary neurons and mouse hnRNPR iCLIP targets from primary motoneurons and NSC34 (motoneuron-line) cell line were downloaded from following publications (Briese et al., 2018; Khudayberdiev et al., 2021). Human and murine genes were converted to fly genes using the DIOPT API tool, and only high-confidence orthologues (DIOPT score >= 8) were retained for comparisons.

#### RBP binding sequence motif discovery

Enriched RBP binding sequence motifs were identified using the Positionally enriched k-mer analysis (PEKA) tool (Kuret et al., 2022). Significant crosslinks (identified from iCount) that are also enriched over SMInput libraries were used as high-confidence thresholded crosslinks (tXn). Immunoprecipitation-recovered crosslinks (*iCount xlsite* output) were used as reference background crosslinks (oXn). Default PEKA settings were used for the analysis with no genome masking. For both Imp and Syp, the 5-mer analysis yielded the highest enrichment scores amongst the 4-8-mers tested. Consensus motifs were plotted using the R package *motifStack* (Ou et al., 2018).

#### Genome interval correlation methods

Relative distances and overlaps (Jaccard indices) between Imp and Syp iCLIP peaks were analysed using the R packages *GenometriCorr* and *valr* (Favorov et al., 2012). For the relative distance metric, the presented correlation index can range from −1 to +1 where the value of −1 indicates perfectly even spacing between binding sites. However, values closer to +1 indicate closer proximity between genomic intervals.

#### Co-evolution analysis

Evolutionary coupling between Imp and Syp binding sites was assessed using the mutual information positional correlations (Weinreb et al., 2016). The pairwise mutual information was calculated as described in (Alfonso-Gonzalez et al., 2023b; Brandman et al., 2012; Martin et al., 2005), and scripts from the Hilgers Lab Zenodo repository were used (Alfonso-Gonzalez et al., 2023a). Briefly, multiple sequence alignment track of 27 *Drosophila* species was downloaded from the UCSC genome browser and filtered using ‘refineMSA()’ function in the Python package *ProDy* (http://prody.csb.pitt.edu/) to keep sequences with 60% gaps (rowocc=0.4) and 98% identity level (seqid=0.98). Nucleotide-level pairwise mutual information scores were normalised using the average product correction (APC) method using ProDy. For each target transcript of Imp and Syp, average mutual information scores were compared between Imp and Syp binding site pairs versus the Imp or Syp binding site columnar average. The comparison was performed in both Imp-to-Syp and Syp-to-Imp directions. For high confidence binding sites, the analysis was limited to Imp and Syp iCLIP peaks that occur in both developmental timepoints were used. Due to computational memory constraints, 5’UTR and 3’UTR portions of the mRNA were computed separately. The resulting mutual information score indicates probability to estimate whether a given nucleotide change will be accompanied by another nucleotide change. For control simulations, shuffled Imp or Syp binding sites for each transcript were generated using ‘bed_shuffle()’ function in the R package *valr*.

#### Hierarchical clustering of RBP binding interactions

To identify groups of transcripts that differentially interact with Imp and Syp across development, we first removed background reads from iCLIP libraries by subtracting average background read counts from non-significant (FDR > 0.01) crosslinks obtained from *iCount peaks* output. For each gene, significant iCLIP reads were then converted to reads per million (RPM), normalised to gene expression levels (transcripts per million, TPM) in the larval brain, and averaged across three biological replicates. We performed *k*-means clustering of normalised iCLIP RPM using the R package *MaSigPro* with following parameters: significance threshold (Q) = 0.01, negative binomial theta = 10, R-squared regression fit threshold = 0.7, clustering method = hierarchical clustering (hclust), k = 6. For plotting purposes only, minimal pseudo-counts were added to genes where no iCLIP read counts were recovered above background.

### Microscopy methods

#### RNA single-molecule fluorescence in situ hybridisation (smFISH)

smFISH was carried out as previously described (L. Yang et al., 2017) with minor modifications. Tissues were dissected in PBS or Schneider’s medium, rinsed once with PBS and fixed with 4% paraformaldehyde (PFA) diluted in PBS/0.1% Triton-X 100 (PBSTx) for 25 min at room temperature (RT). Samples were rinsed briefly in PBSTx and then further permeabilised with PBSTx twice for 20 min each. Samples were pre-hybridised in smFISH wash solution (10% formamide, 2xSSC) for 30 min at 37°C and hybridised in smFISH hybridisation solution (10% formamide, 2xSSC, 10% dextran sulphate) with probes diluted to 250nM final concentrations overnight at 37°C. Samples were rinsed briefly in smFISH wash solution and then washed in smFISH wash solution twice for 30mins each at 37°C. Finally, samples were washed in 2xSSC for 10mins at RT and mounted on slides using either Vectashield (Vector labs) or Slowfade Diamond (ThermoFisher) anti-fade mounting media. Slides were either imaged immediately or stored at 4°C. Counterstains were included in the wash solution: DAPI (1μg/ml), fluorophore-conjugated phalloidin (1:100, ThermoFisher). smFISH probe sequences used in this study are available in Supplementary Table 5.

#### smFISH probe synthesis

Candidate smFISH probe sequences were generated using the Stellaris Probe Designer version 4.2 (https://www.biosearchtech.com/stellaris-designer) with the following parameters: ‘masking level, 5; oligo length, 20nt; minimum spacing length, 3nt’. Oligonucleotides were singly labelled with ATTO488, ATTO633, Alexa Fluor 647, or Cy3 at the 3’ ends according to a published protocol (Gaspar et al., 2017). Dye-labelled oligos were purified using Oligo Clean and Concentrator Kit (Zymo Research), and the probe concentrations were normalised to 25μM using nanodrop measurements. All probe sets used in this study had a degree of labelling > 0.96.

#### Immunofluorescence

Tissues were dissected and fixed as with smFISH procedure. After permeabilisation, samples were blocked in blocking solution (Licor Odyssey blocking buffer supplemented with 0.1% Tween-20) for 30-60min at RT. Then, samples were incubated with primary antibodies diluted in blocking solution. Samples were probed overnight at 4°C and were washed three times in PBS/0.1% Tween-20 (PBSTw) for 10 min each at room temperature and incubated with fluorescent secondary antibodies (1:500) diluted in blocking solution overnight at 4°C. After further three washes in PBSTw, cells were mounted in an anti-fade mounting medium. Antibodies used in this study are listed in the Resource Table. For combined smFISH and immunofluorescence, antibody staining was carried out sequentially after the smFISH protocol. Additionally, the blocking solution was pre-treated with 1:25 RNASecure (Invitrogen) for 30min at 50°C and supplemented with 2mM RVC to prevent RNase activity.

#### Puromycin incorporation assay

Dissected L3 larval brains were washed briefly in brain culture media (80% Schneider’s medium, 20% fetal bovine serum, 100 µg/ml insulin) and incubated in brain culture media with or without 5 µg/ml puromycin. Tissues were incubated *ex vivo* for 1 hour at room temperature with shaking at 100 RPM. For control experiments, 100 µg/ml cycloheximide (CHX) treatment was added at the same time with puromycin. Incorporated puromycin was visualised using anti-puromycin (Sigma, ZMS1016) via immunofluorescence. Only the top layer of central brain NB lineages was imaged to ensure an even degree of puromycin penetration between tissues.

#### Image acquisition and handling

Mounted specimens were imaged on an Olympus SpinSR10 spinning disk confocal system equipped with Prime BSI camera or on an Olympus FV3000 Inverted Laser Scanning Microscope. Objectives used were ×20 dry (0.8 NA, UPLXAPO20X), ×60 silicone oil (1.3 NA, UPLSAPO60XS2), or ×100 oil (1.45 NA, UPLXAPO100XO). Image voxel sizes were 0.55 × 0.55 × 2 μm (x:y:z) with the ×20 objective and 0.11 × 0.11 × 0.2 μm (x:y:z) with the ×60 and ×100 objectives. Microscopes were driven using Olympus cellSens Dimension or FV313S software. Images were uploaded and stored in the University of Oxford OMERO server (Allan et al., 2012), and OMERO.figure was used to generate image visualisations.

#### Fluorescence intensity quantification

Immunofluorescence images were background subtracted using the rolling ball subtraction method (radius = 50-150 px) in ImageJ. Antibody stain was quantified by integrating fluorescence intensity across the z-stacks of region of interest (ROI) divided by the volume to obtain signal density. Camera background signals (area with no tissue) were subtracted from the signal density, and the resulting values were normalised to control conditions.

#### smFISH image quantification

smFISH images were quantified as described previously (Lee et al., 2022) using a custom Python pipeline using *Bigfish* (Imbert et al., 2021), *Sci-kit image* and *Numpy* libraries. Briefly, exonic smFISH channels were background subtracted with the *skimage.white_tophat* algorithm (radius = 5 px) and Laplacian of Gaussian (LoG) filtered with a theoretical point spread function based on microscope acquisition settings. Thresholds for spot detection were set manually based on the intensity of LoG filtered spots, and reference single-molecule images were obtained using the *build_reference_spot()* function iterating over the entire image dataset in Bigfish. Intranuclear transcription sites were localised using intronic smFISH channels and corresponding exonic channels were used to resolve the number of nascent RNA using the *decompose_cluster()* function in Bigfish.

#### RNA half-life calculation

RNA half-lives were calculated using the *TransQuant* steady-state methodology (Halpern and Itzkovitz, 2016). A MATLAB-to-R translated version of TransQuant script was used to obtain the probe library weighting factor, which accounts for the positional information of each oligonucleotide against the target transcript. Transcription rate, decay rate, and half-lives were calculated using the equations listed below. An elongation rate of 90 kb per hour was used for the calculation (Samuels et al., 2020b). For *Eip93F*, chromosome fraction was 2. For *jim* and *Ldh*, the proportion of cells positive for intronic smFISH signal per field of view was used as chromosome fraction.

● Transcription rate (mRNA*hour^-1^) = ((nascent transcript number/weighting factor)* elongation rate)/gene length
● Decay rate (hour^-1^) = (chromosome fraction x transcription rate x number of chromosome copies)/transcripts in the cell
● Half-life (min) = (ln(2)/decay rate) * 60

## Statistics and data presentation

All statistical analyses were performed using the R package *rstatix*. Data normality was assessed using the Shapiro-Wilk test. Comparisons were performed either using the parametric tests, t-test and ANOVA-test or non-parametric tests Mann-Whitney U or Dunnett’s test. Where applicable, *p*-values were adjusted using Bonferroni’s method for multiple comparisons. For data wrangling, the *tidyverse suite* of packages were used in the RStudio environment, while *Numpy* and *Pandas* Python libraries were used in the Jupyter notebook environment. The R packages *ggplot2*, *ggbeeswarm*, *scales*, and *patchwork* were used to create the presented visualisations. Further visual annotations were made in Affinity Designer (Serif).

## Data availability

Raw FASTQ and processed iCLIP sequencing data files are deposited to Gene Expression Omnibus (GEO). Reviewers’ token for read-only access is available upon request. All other source data are available from the Github repository: https://github.com/jefflee1103/Lee2024_Imp-Syp-iCLIP.

## Author contributions

J.Y.L. - Conceptualization, Data curation, Formal analysis, Investigation, Visualization, Methodology, Project administration, Writing - original draft, Writing - review and editing. N.H. - Data curation, Formal analysis, Investigation, Methodology. T.J.S. - Conceptualization, Data curation, Investigation.

I.D. - Conceptualization, Supervision, Methodology, Resources, Funding acquisition, Writing - review and editing.

## Funding

Wellcome Trust (Investigator Award 209412/Z/17/Z) - J.Y.L., N.H., T.J.S., I.D. University of Oxford (Medical Sciences Graduate Studentship) - J.Y.L. Wellcome Trust (Four-Year PhD Studentship 105363/Z/14/Z) - T.J.S.

## Acknowledgement

We are grateful to the University of Oxford Micron imaging facility for help with advanced microscopy. We thank Amanda Williams for the operation of the Next-generation sequencing facility. Fly stocks and antibodies were kindly gifted by Chris Doe and Alberto Baena-Lopez. We are grateful to Alfredo Castello, Francesca Robertson, Mary Kay Thompson, Joshua Titlow, and Aino Jarvelin for their advice on the experimental design, and to Richard Parton, Ana Palanca and Darragh Ennis for their advice on microscopy and image handling.

## Conflicts of interest

No competing interests declared.

---

**Figure S1.**
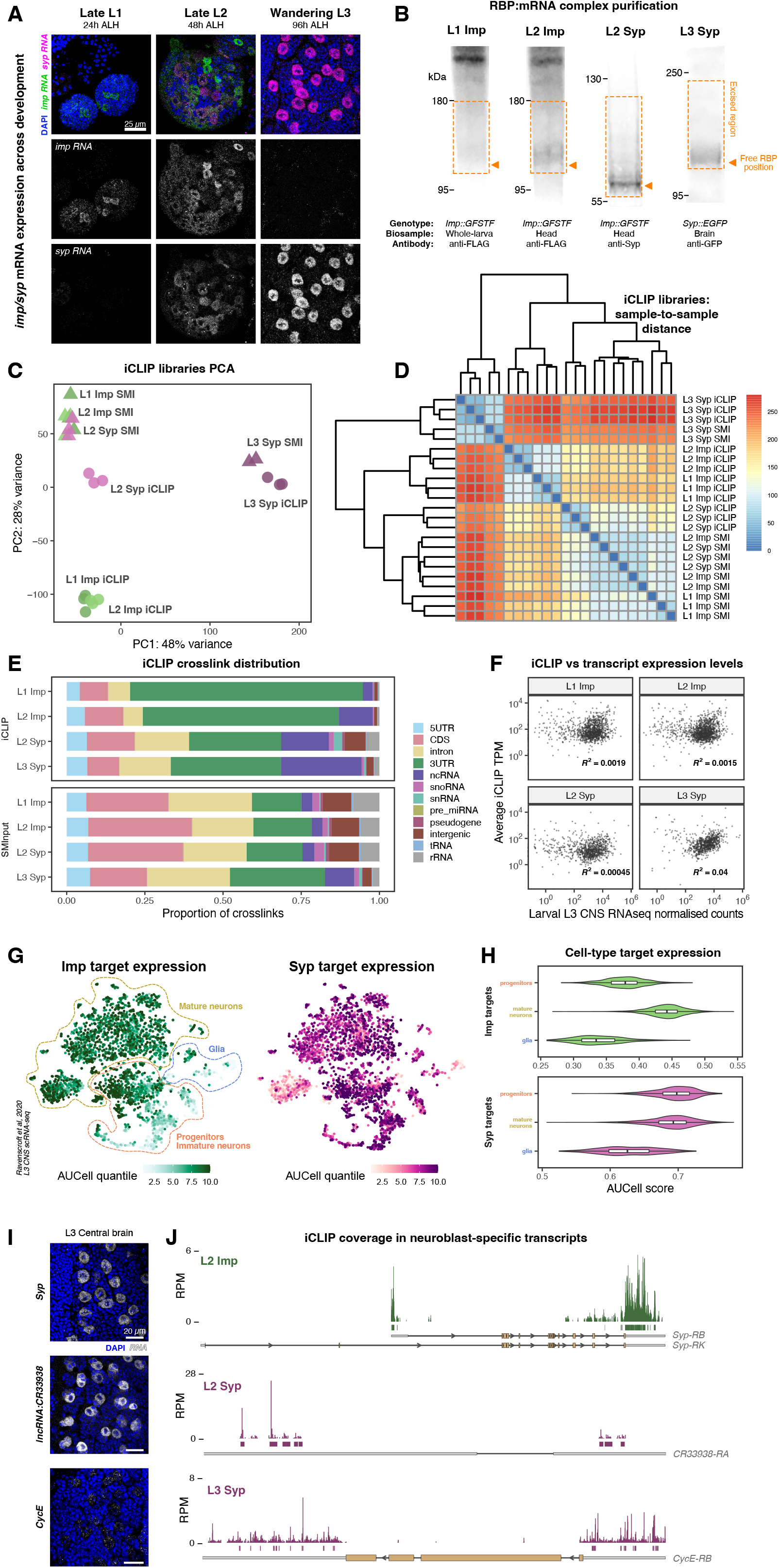
Identification of *in vivo* RNA targets of Imp and Syp in larval brains, related to Figure 1. **(A)** Temporal expression pattern of *imp* and *syp* RNA in *Drosophila* larval nervous system, visualised using single-molecule fluorescence *in situ* hybridisation (smFISH). **(B)** Infrared blot of RNA-binding protein (RBP):RNA complexes purified via immunoprecipitation. Orange regions indicate the excised molecular weight range used for the iCLIP library preparation. Orange arrows indicate expected molecular weight of unbound Imp or Syp. **(C)** Principal component analysis (PCA) of iCLIP libraries based on normalised crosslink counts per gene. Each iCLIP library is denoted with a different colour, and size-matched input (SMInput) libraries are indicated with a different shape. **(D)** Heatmap of a hierarchical clustering of iCLIP and SMInput libraries. The values indicate sample-to-sample distance calculated from transformed count matrices. **(E)** Relative proportion of iCLIP and SMInput crosslinks for each library grouped by transcript biotypes. Protein coding genes were further subdivided into 5’UTR, CDS, 3’UTR, and intron. **(F)** Correlation of iCLIP reads (normalised to transcripts per million, TPM) versus transcript abundance obtained from Wildtype L3 brains (Samuels et al., 2020a) per gene. Each symbol indicates average values from three replicates, and R^2^ values are shown for each iCLIP library. **(G)** Area Under Curve Cell (AUCell) scores of Imp and Syp targets across the L3 brain single-cell RNA-seq atlas (Ravenscroft et al., 2020). Broad progenitors/immature neurons, mature neurons and glial cell types identified from the original publication are indicated. **(H)** Comparison of AUCell scores of Imp and Syp targets between progenitors, mature neurons or glia cell-types. **(I)** RNA expression patterns of neuroblast (NB) specific transcripts, *Syp*, *lncRNA:CR33938* and *CycE*, visualised using smFISH in the L3 brain. **(J)** Imp and Syp iCLIP coverage in NB-specific transcripts.

**Figure S2.**
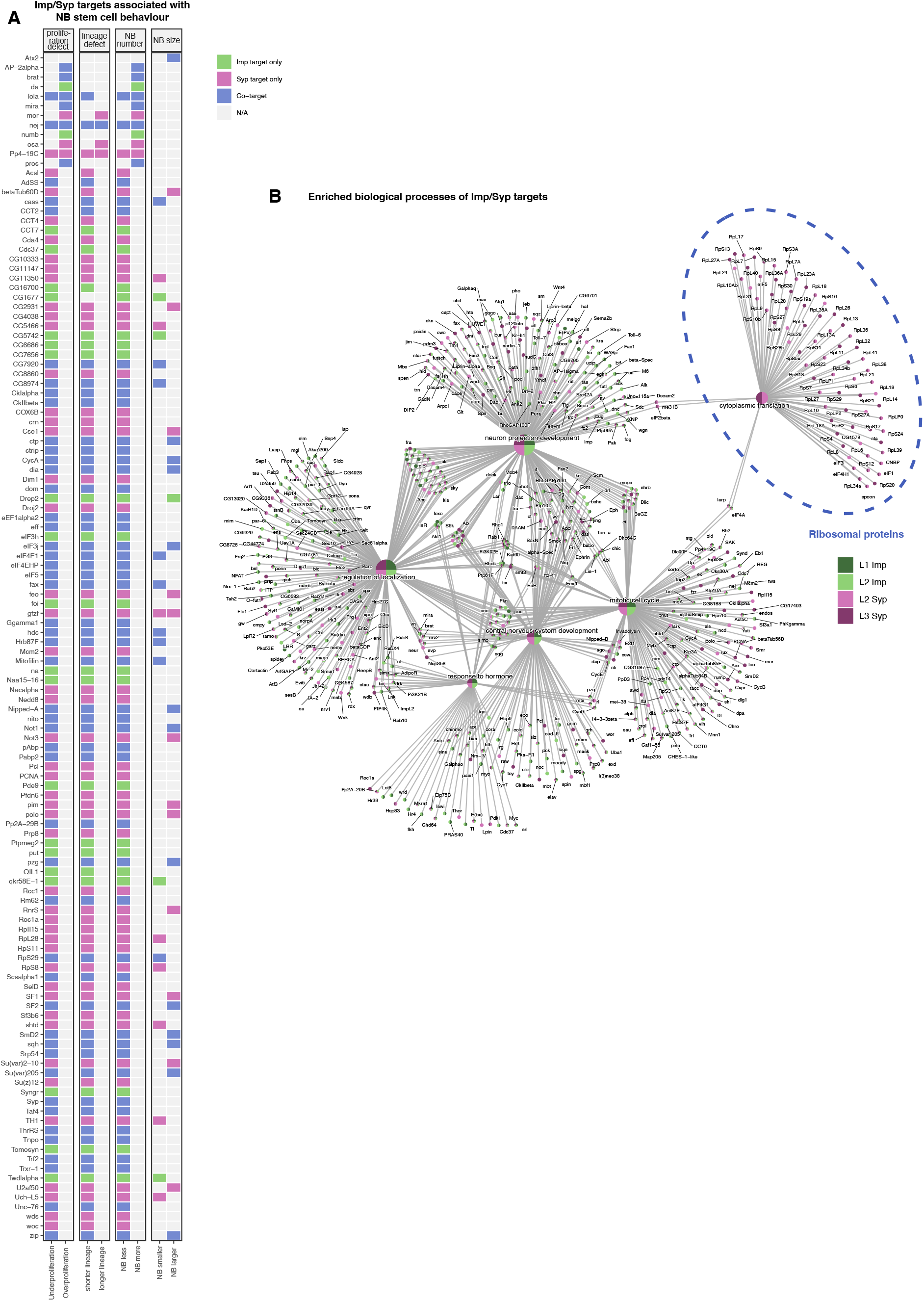
The analysis of Imp and Syp target functions in the neuroblast lineage, related to Figure 2. **(A)** Imp and Syp targets that regulate NB size, NB number, NB lineage length, or neural progenitor proliferation, as identified from a genome-wide RNAi screen (Neumuller et al., 2011). Only the genes that are Imp or Syp targets are shown. **(B)** Network plot of top enriched GO biological process terms and associated genes. The shades of green and magenta colours represent Imp or Syp iCLIP target status. A group of transcripts encoding ribosomal proteins is indicated in blue, which exclusively interacts with Syp.

**Figure S3.**
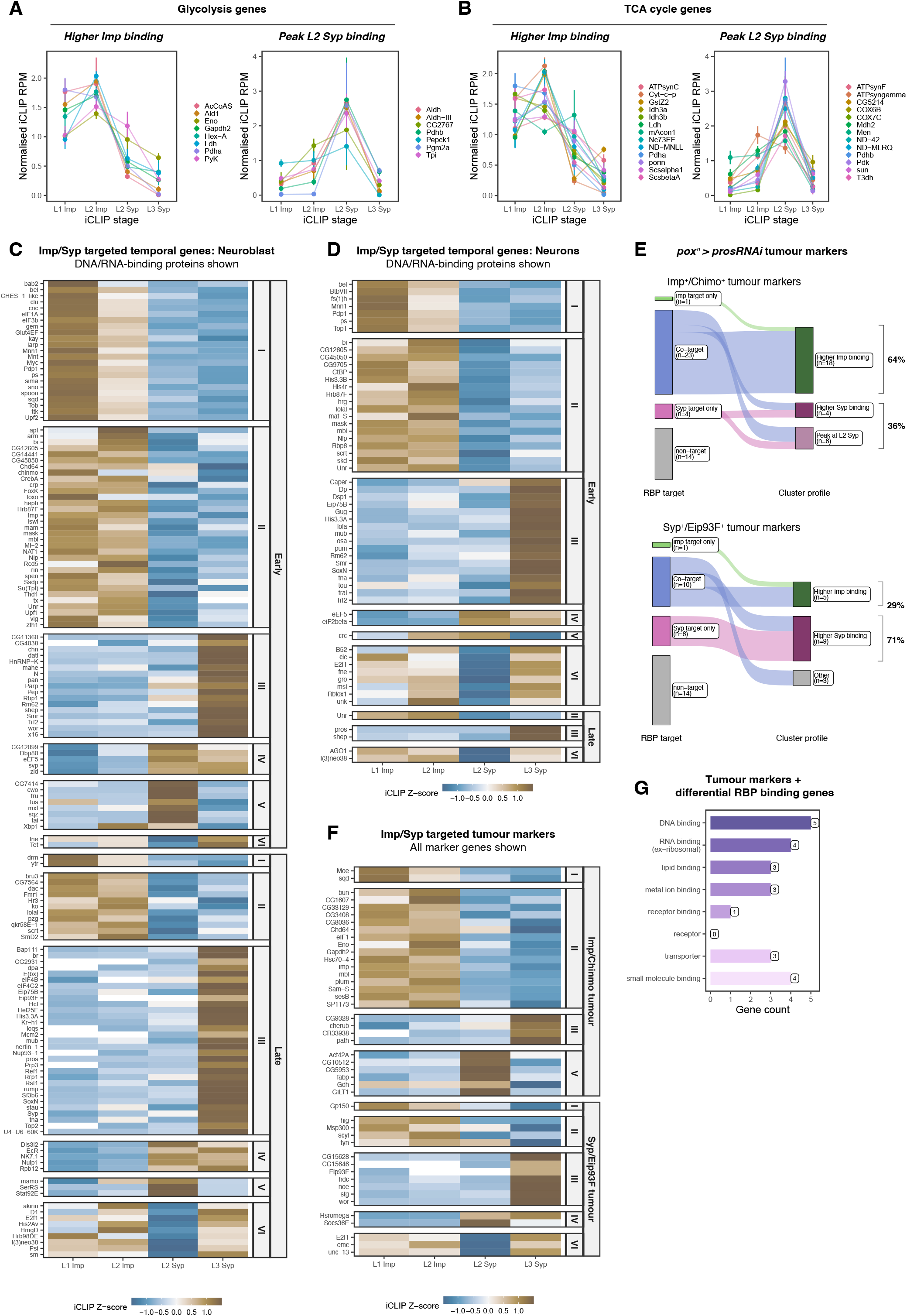
Temporally expressed transcripts interact dynamically with Imp and Syp, related to Figure 3. (**A-B**) Transcripts encoding regulators of energy metabolism involved in (**A**) glycolysis or (**B**) TCA cycle that show ‘High Imp binding’ or ‘Peak L2 Syp binding’ interaction patterns. (**C-D**) Heatmaps showing scaled iCLIP scores of temporally regulated factors in (**C**) NB or (**D**) immature/mature neurons grouped by their *k*-means Cluster membership. Only the transcripts encoding DNA or RNA-binding proteins are shown. **(E)** Sankey plot of hierarchical tumour markers (Imp^+^/Chinmo^+^ versus Syp^+^/Eip93F^+^) that are desbried to dynamically interact with Imp and Syp. Tumour markers were taken from pox^n^ > *pros* RNAi scRNA-seq dataset (Genovese et al., 2019). **(F)** Heatmaps showing scaled iCLIP scores of Imp^+^/Chinmo^+^ and Syp^+^/Eip93F^+^ tumour markers and their *k*-means Cluster membership. **(G)** Enriched molecular function of Imp^+^/Chinmo^+^ and Syp^+^/Eip93F^+^ tumour markers that dynamically interact with Imp and Syp. The terms represent GOSlim terms following the FlyBase molecular function ribbon classification.

**Figure S4.**
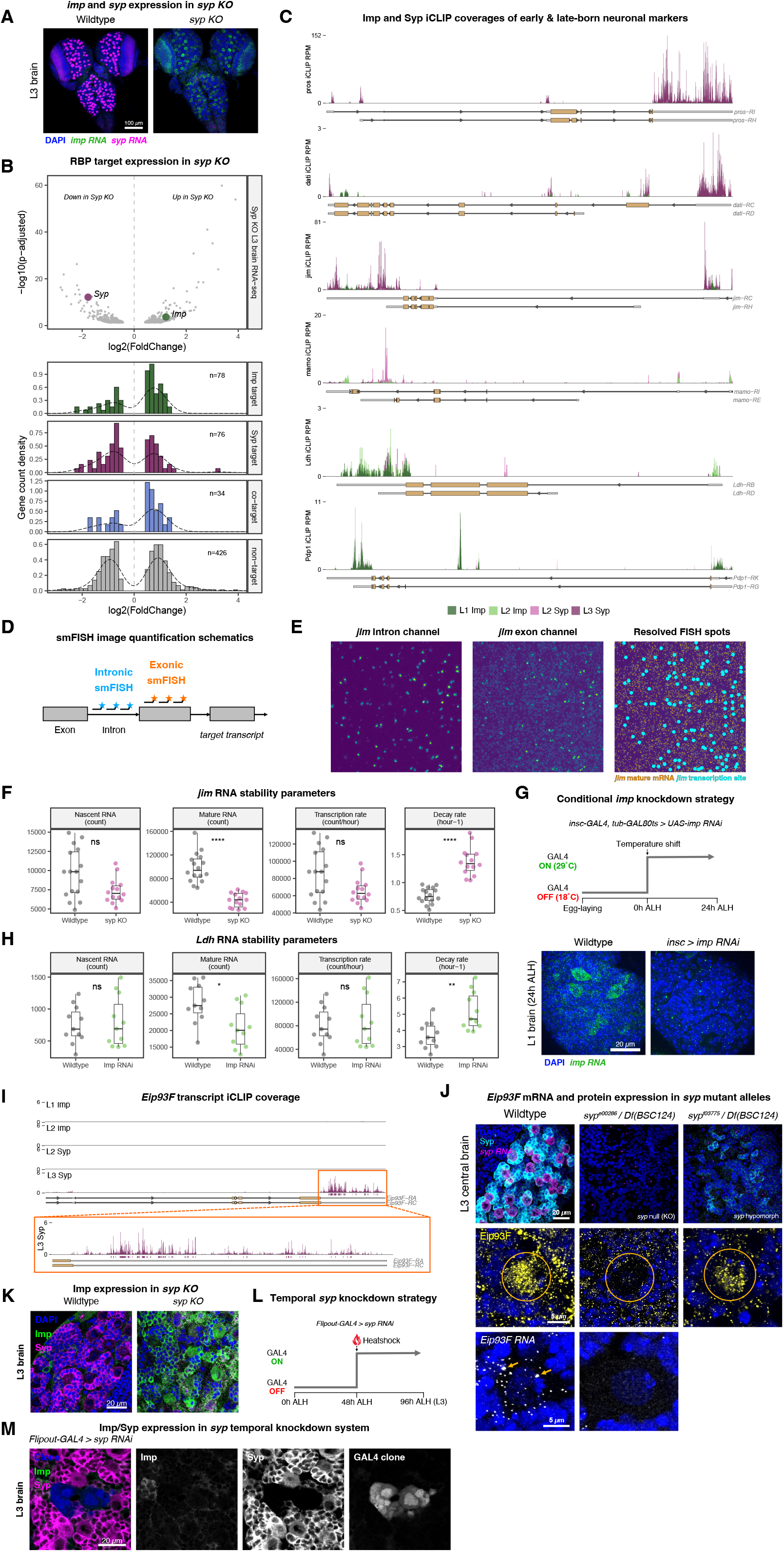
Imp and Syp post-transcriptionally regulate downstream targets, related to Figure 4. **(A)** Expression of *imp* and *syp* mRNA in the Wildtype and *syp KO* L3 brains. Note the sustained expression of *imp* in the L3 stage upon *syp KO*. RNA expression of both *imp* and *syp* are enriched in NBs. **(B)** Imp and Syp targets that are differentially expressed in L3 *syp KO* brains. In *syp KO* brains, *syp* is downregulated (magenta symbol) and *imp* is upregulated (green symbol). The histograms show the distribution of log2 fold change in transcript abundance for Imp, Syp or co-targets. **(C)** iCLIP coverages of Imp and Syp across early and late immature neuron marker transcripts. **(D)** smFISH strategy to calculate steady-state RNA stability using exon and intron probes. **(E)** Example images of intron and exon smFISH channels of *jim* mRNA in the larval central brain. Both channels were used to count mature and nascent transcripts using BigFISH (see Methods). Viridis look-up table is used. **(F)** Quantification of *jim* RNA stability parameters. (n=3). **p<0.0001. **(G)** Strategy to knockdown *imp* in a temporal manner using *insc-GAL4* and *tub-GAL80ts*. The smFISH image of L1 *imp* knockdown brain shows presence of *imp* transcription (bright foci) but lack of mature transcript accumulation. **(H)** Quantification of *Ldh* RNA stability parameters. (n=2). **p<0.01. **(I)** iCLIP coverages of Imp and Syp on *Eip93F* mRNA. There are no significant Imp and L2 Syp binding sites on *Eip93F*. **(J)** Expression pattern of *Eip93F* RNA and protein in Wildtype, *syp KO* (*syp* null allele over deficiency), and *syp hypomorph* (*syp* hypomorphic allele over deficiency). Yellow arrows indicate nuclear transcription sites of *Eip93F*, which is absent in *syp KO* NBs. **(K)** Expression pattern of Imp and Syp proteins in Wildtype or *syp KO* L3 brains. Central brain regions are shown. **(L)** A strategy to temporally knockdown *syp* using an inducible *flip-out GAL4* system. **(M)** Expression patterns of Imp and Syp in the temporal *flip-out GAL4 > syp* RNAi NB clones. Note that this system allows specific knockdown of Syp without Imp overexpression.

**Figure S5.**
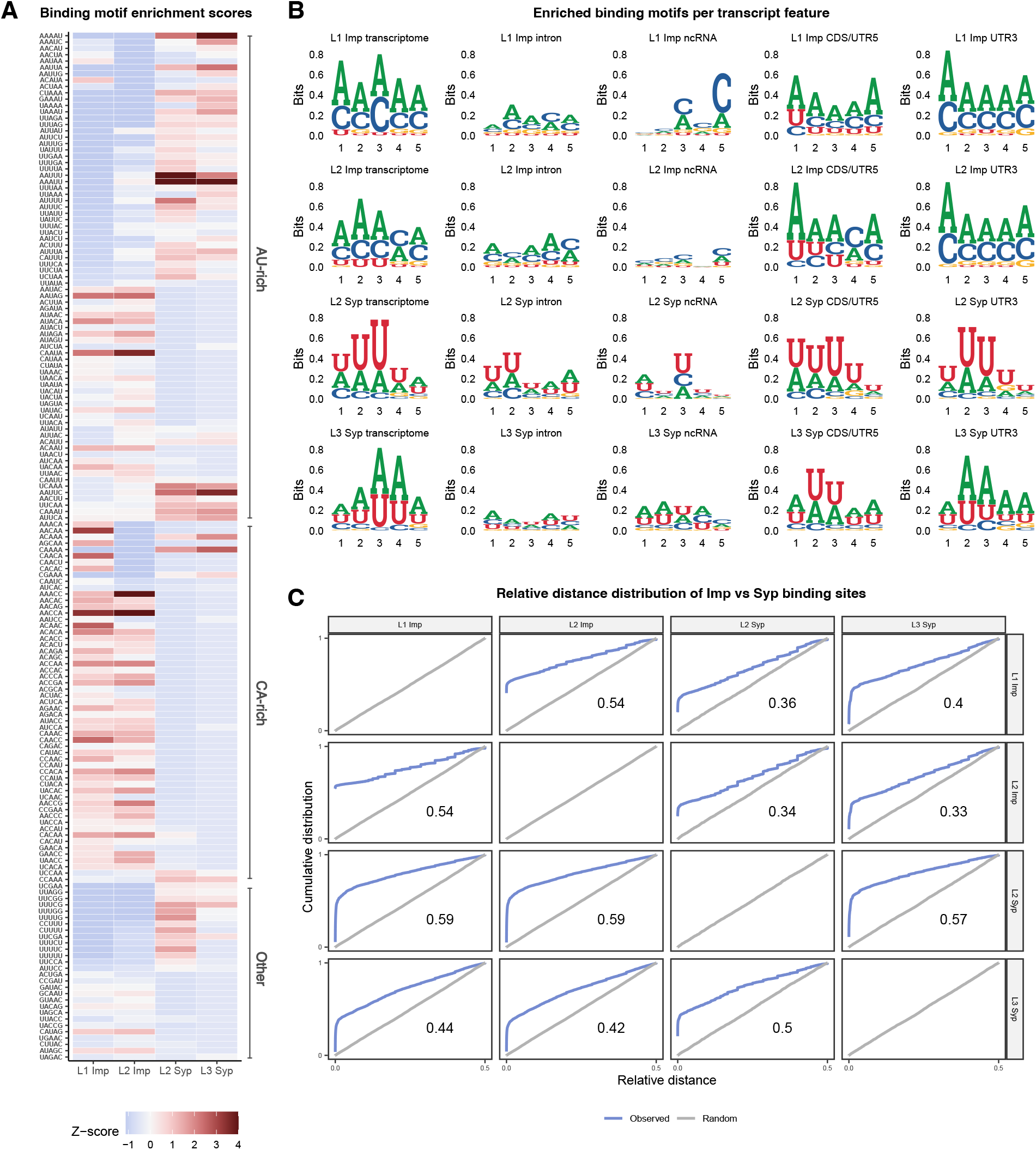
Imp and Syp binding sites poorly overlap, except in temporally expressed transcripts, related to Figure 5. **(A)** Heatmap of RNA sequence motif enrichment scores (scaled) for each iCLIP library. Only the top enriched 5-mers are shown. The plot is subdivided into AU-rich, CA-rich, and other enriched sequence motifs. **(B)** Enriched RNA binding sequence motifs of Imp and Syp across different developmental stages and transcript features. **(C)** Relative distance metrics between Imp and Syp iCLIP binding sites. Cumulative plots that compare observed (empirical) relative distances versus randomly simulated binding sites are shown. Transcriptome-wide relative distance metrics were calculated using the *genometricorr* R package. The correlation index can range from −1 to +1 where the value of −1 indicates perfectly even spacing between binding sites, while values closer to +1 indicate closer proximity between genomic intervals.

**Figure S6.**
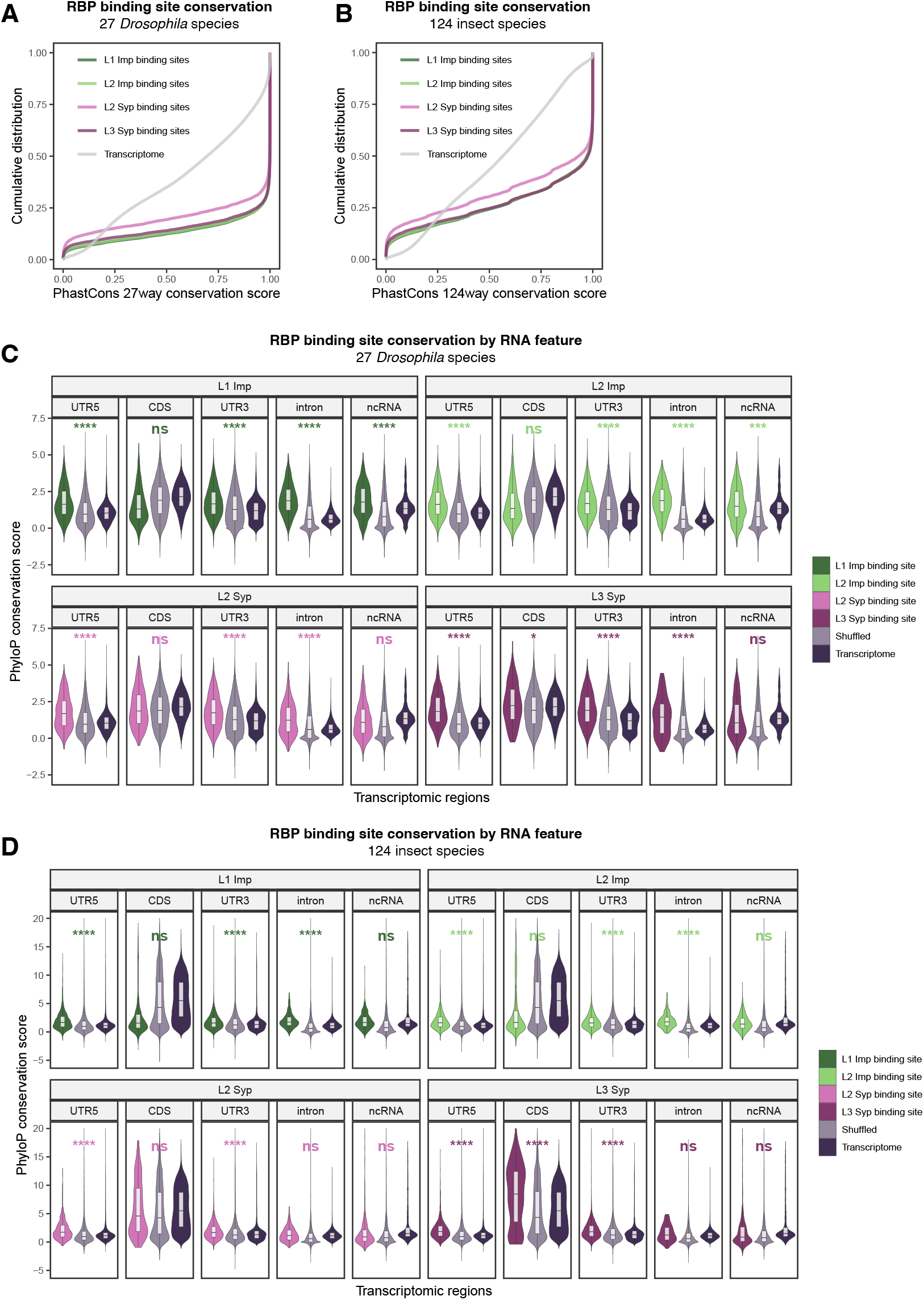
Imp and Syp binding sites display evolutionary linkage, related to Figure 6. (**A-B**) Comparison of PhastCons sequence conservation score of Imp and Syp binding sites from each iCLIP library versus the transcriptome average using (**A**) the 27-way *Drosophila* or (**B**) 124-way insect PhastCons track (downloaded from the UCSC genome browser). (**C-D**) Comparison of PhyloP sequence conservation score of Imp and Syp binding sites from each iCLIP library versus the brain transcriptome and the shuffled iCLIP binding sites per gene per transcript features using (**C**) the 27-way *Drosophila* or (**D**) 124-way insect PhyloP score matrices. For shuffled binding sites, the simulation was iterated 10 times per query. *p<0.05, ***p<0.001, ****p<0.0001.

## Supplementary file legend

**Supplementary table 1: Summary of Imp and Syp iCLIP targets.**

exp_string: iCLIP RNA-binding protein and developmental stage; gene_id: FlyBase gene ID (Ensembl release 99); gene_name: FlyBase gene name; gene_biotype; gene biological type; n_binding_site: number of significant RBP binding sites; binding_feature: iCLIP binding site transcript features; l2fc_xlsite: log2FoldChange of crosslinks compared to SMInput across the entire gene body; padj_xlsite: adjusted p-value of crosslinks compared to SMInput across the entire gene body; l2fc_bsmax: log2Foldchange of the most enriched binding site compared to SMInput; padj_bsmin: minimum adjusted p-value within binding sites compared to SMInput; feature_pct: percentage of iCLIP binding sites overlapping with each transcript feature; avg_iCLIP_tpm: average crosslink transcripts per million; human_homologs: high confidence human homologs (DIOPT score > 7); mammalian_imp_target: conserved target with mammalian IMP1-3 in human pluripotent stem cells (Conway et al., 2016); mammalian_syp_target: conserved target with mammalian SYNCRIP or HNRNPR in rodent neurons (Briese et al., 2018; Khudayberdiev et al., 2021).

**Supplementary table 2: GO enrichment analysis of Imp and Syp iCLIP targets.**

GO_type: Gene Ontology (GO) class; exp_string: iCLIP RNA-binding protein and developmental stage; GO_id: GO accession number; Description: GO ontology term; GeneRatio: ratio of genes annotated with each GO term in the foreground set; BgRatio: ratio of genes annotated with each GO term in the brain transcriptome background set; p.adjust: adjusted p-value of the enrichment hypergeometric test; gene_names: annotated genes in the foreground set for each GO term; l2fc: log2FoldChange enrichment of each GO term.

**Supplementary table 3: Differential occupancy Imp and Syp targets.**

gene_id: FlyBase gene ID; gene_name: FlyBase gene name; imp_target: L1_Imp or L2_Imp iCLIP target; syp_target: L2_Syp or L3_Syp iCLIP target; cluster_id: differential occupancy cluster grouping; cluster_description: differential occupancy cluster description; sypKO_l2fc: log2FoldChange of transcript expression comparing sypKO versus wildtype L3 brains; sypKO_padj: adjusted p-value of sypKO_log2fc.

**Supplementary table 4: Co-evolution analysis output between Imp and Syp binding sites.**

gene_id: FlyBase gene ID; gene_name: FlyBase gene name; row_coverage: total nucleotide coverage of query binding sites; row_mean: mean mutual information score across total nucleotide coverage of query binding sites; row_median: median mutual information score across total nucleotide coverage of query binding sites; region_coverage: nucleotide coverage of Imp and Syp binding sites; region_mean: mean mutual information score across pairwise Imp and Syp binding sites; region_median: median mutual information score across pairwise Imp and Syp binding sites; tstatic: t-test statistics; pvalue: t-test p-value; comparison_mode: comparison direction from query to reference, either Imp-to-Syp or Syp-to-Imp; feature: comparison transcript feature, either UTR5 or UTR3.

**Supplementary table 5: smFISH probe sequences used in this study.**

smFISH probe sequences were designed using the online Stellaris probe designer tool (https://www.biosearchtech.com/stellaris-designer) and ordered as unmodified oligonucleotides from IDT (see Methods).

## Notes

### Competing Interest Statement

The authors have declared no competing interest.

